# Multi-Agent Reinforcement Learning-based Adaptive Sampling for Conformational Sampling of Proteins

**DOI:** 10.1101/2022.05.31.494208

**Authors:** Diego E. Kleiman, Diwakar Shukla

## Abstract

Machine Learning is increasingly applied to improve the efficiency and accuracy of Molecular Dynamics (MD) simulations. Although the growth of distributed computer clusters has allowed researchers to obtain higher amounts of data, unbiased MD simulations have difficulty sampling rare states, even under massively parallel adaptive sampling schemes. To address this issue, several algorithms inspired by reinforcement learning (RL) have arisen to promote exploration of the slow collective variables (CVs) of complex systems. Nonetheless, most of these algorithms are not well-suited to leverage the information gained by simultaneously sampling a system from different initial states (e.g., a protein in different conformations associated with distinct functional states). To fill this gap, we propose two algorithms inspired by multi-agent RL that extend the functionality of closely-related techniques (REAP and TSLC) to situations where the sampling can be accelerated by learning from different regions of the energy landscape through coordinated agents. Essentially, the algorithms work by remembering which agent discovered each conformation and sharing this information with others at the action-space discretization step. A *stakes function* is introduced to modulate how different agents sense rewards from discovered states of the system. The consequences are threefold: (i) agents learn to prioritize CVs using only relevant data, (ii) redundant exploration is reduced, and (iii) agents that obtain higher stakes are assigned more actions. We compare our algorithm with other adaptive sampling techniques (Least Counts, REAP, TSLC, and AdaptiveBandit) to show and rationalize the gain in performance.

## 1 Introduction

Molecular dynamics (MD) simulations are a well-established computational method in chemistry, condensed matter physics, materials science, and biology. They are useful to probe properties of atomic-scale systems at time scales and levels of resolution rarely accessible by experimental techniques.^1^ Despite the significant improvements in the performance of hardware and software utilized to run MD simulations, sampling conformational changes in large systems remains challenging in common practice.^2^ This is due to the onerous computational cost of running MD simulations. In biological applications, most modern studies are restricted to sampling systems for a few microseconds only. However, cellular processes of interest occur at slower time scales, such as protein folding,^3^ activation of signalling proteins,^4,5^ transport across transmembrane proteins,^6–8^ and ligand binding.^9,10^

To alleviate this problem, numerous enhanced sampling methods have been proposed.^11^ The shared goal of these techniques is to reduce the required run time to obtain adequate sampling of the system at hand. Nonetheless, different techniques are usually developed with different applications in mind, and thus the best choice of method may vary depending on the goal of a study. Enhanced sampling methods can be roughly divided into two categories depending on their requirement of collective variables (CVs). CVs are functions of one or more degrees of freedom of the system and they are useful because they can describe state transitions in a reduced dimensional space.^12^ Methods that require the user to define appropriate CVs use this information to bias the sampling along these coordinates to accelerate the exploration of states of interest. A subset of this class of methods (e.g., metadynamics, umbrella sampling) achieve this by biasing the underlying potential along the CVs.^13,14^ A different subset of CV-based methods, which are the focus of this study, statistically bias the sampling along user-defined coordinates or bins by adaptively selecting new simulation starting points (e.g., Least Counts adaptive sampling, weighted ensemble MD).^15–17^ These methods are particularly well-suited to exploit the power of large clusters through parallel simulations.^2,17,18^ It must be noted that some adaptive sampling methods are not dependent on reaction coordinates.^19,20^ The second class of enhanced sampling methods alter the systems Hamiltonian to enhance exploration across all degrees of freedom (e.g., accelerated MD, REMD-SSA).^21,22^

Since MD simulations typically seek to explore relevant regions of the systems phase space, a natural trade off that often occurs is termed the exploration-exploitation dilemma.^23^ The dilemma is whether we should devote computational resources to sample already explored areas of the phase space to improve the estimate of a metric of interest (e.g., the free energy of a proteins metastable state) or explore new areas of the space to find new relevant regions (e.g., a rare state transition). Diverse forms of this dilemma occur in many areas of science and have been addressed particularly by a branch of machine learning (ML) denominated reinforcement learning (RL).^24^ In RL, the optimization problem is formulated in terms of reward maximization rather than loss minimization.^25^ Developments in ML methods have informed the workflows used by MD simulation practitioners. ML-based techniques are now used to both conduct MD simulations and analyze the resulting data.^26,27^ In this way, several enhanced sampling techniques based on RL principles or algorithms have arisen (e.g., FAST,^23^ REAP,^28^ TSMD,^29^ AdaptiveBandit^30^ and TALOS^31^).

Since the thermodynamic ensembles sampled by MD simulations usually follow a Boltzmann distribution, the probability of sampling a state decays exponentially with its energy. For this reason, exploration of the rare transitions is often more computationally challenging than the sampling of known metastable states. Therefore, RL-based enhanced sampling methods tend to focus on encouraging the exploration of the energy landscape under the assumption that researchers can later extend sampling of discovered states to reduce the uncertainty of their measurements.

RL-based adaptive sampling (REAP) is a technique that enhances the exploration of the energy landscape along user-defined CVs. Unlike other similar techniques, it utilizes a reward function where each CV is weighted based on knowledge from the data collected so far. These weights are iteratively updated, so that the user can interpret which CV is more relevant at a given point of the simulation. In general the REAP algorithm can be summarized as follows: a set of MD trajectories is first clustered to discretize and reduce the size of the set of conformations from which new simulations can be run (action space discretization). Clusters with the lowest number of members are selected as candidates. Then, the weights of the CVs are updated such that the cumulative rewards from the candidates is maximized. Finally, new simulations are spawned from the conformations with highest rewards. This cycle is repeated until the system reaches a desired final state.^28^

REAP was intended to explore free energy landscapes where a few orthogonal CVs can describe the relevant transitions. Biologically relevant examples of such landscapes include kinase activation,^4,32,33^ transmembrane transport,^34^ and protein folding.^3^ In those cases, the system has to diffuse across orthogonal CVs to reach the final state. Here, REAP can effectively push the system from one state to another by learning which CV must be prioritized.^28^

However, the functionality of REAP is limited by the assumption that new regions of the landscape will tend to show new extrema along the prioritized CVs. A simple example where such assumption is violated is when MD practitioners start sampling a system from different states. For example, a transporter may be studied via MD simulations through parallel runs from its inward facing and outward facing states. The REAP algorithm would not be suitable in this situation because it preferentially selects conformations that show extreme values of the CVs. However, it is possible that unexplored regions of the conformational landscape that are crucial for the transition between states are contained within the range of the CVs (Figure 1c). In such cases, even the simpler Least Counts (LC) sampling scheme might show better performance at capturing this transition (Figure 1a).^16,35^ Moreover, we can imagine a case where a researcher starts independent simulations from different states of a system but does not combine the resulting data until sampling is deemed appropriate. In this case, the risk is that regions of the landscape that were already discovered are redundantly sampled, thus wasting computational resources.

**Figure 1:**
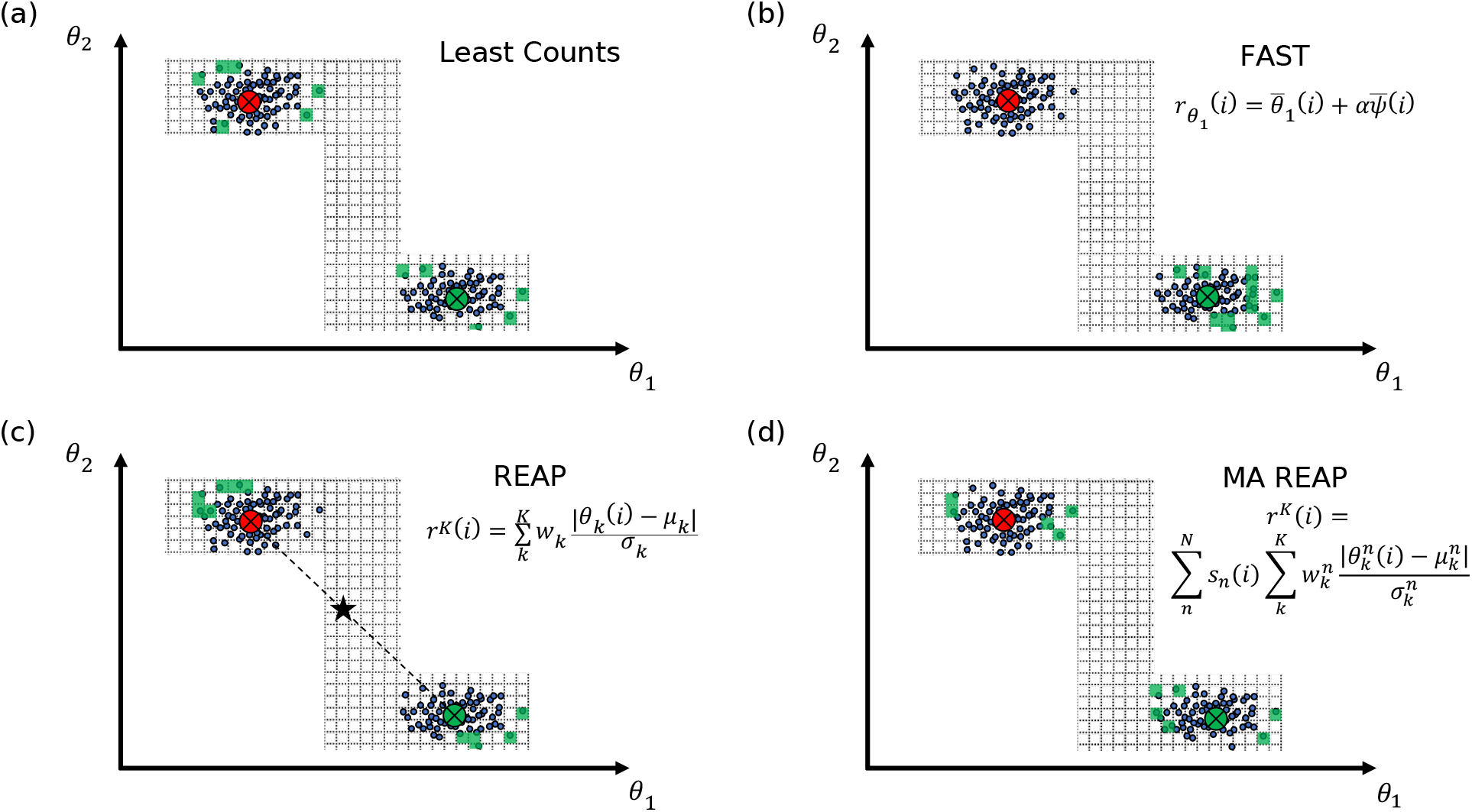
Diagrams showing the different behavior of adaptive sampling algorithms in a hypothetical landscape. The red and green X’s mark the starting structures for the sampling. The states are divided by a grid, rather than clusters. Green grid cells represent states that would be rewarding for each algorithm. The star in (c) represents the approximate location of the mean of the data distribution. (a) Least Counts selects restarting states that contain the fewest observed frames. (b) FAST employs a binomial reward function that biases the choice of restarting states towards the states with maximum (or minimum) values of a single CV. The undirected term favors exploration by rewarding states with low frame counts. (c) REAP assigns rewards based on the distance of conformations to the average of the population. If sampling is started in two different states, pooling all the data together may result in a reward function that only favors conformations that exhibit extreme values of the CVs, rather than conformations that approach a rare or intermediate state. (d) Multi-agent REAP incorporates multiple agents that compartmentalize their data, allowing them to independently sample along high-variance CVs and sharing data only when in proximity to another agent’s discovered states.

In order to provide a sampling scheme that incorporates the benefits of REAP and is better suited to simultaneously harness data from different regions of a free energy landscape, we turned to multi-agent reinforcement learning (MARL) to develop multi-agent (MA) REAP. For the remainder of the paper, the terms REAP and single-agent (SA) REAP will be used interchangeably and in contrast to the multi-agent implementation. An agent is defined as a learning system that must find an optimal function (the policy) to map agent states (e.g., set of discovered conformations) into actions (e.g., which conformations to use to launch new trajectories). In most MD literature drawing on MARL terminology, there is a tendency to equate an agent with a single trajectory (multiple walker metadynamics,^36^ TALOS^31^). However, such correspondence is not theoretically required and in this study we think of an agent as a mathematical model that manages a set of simulations on behalf of the researcher.

In comparison to the baseline REAP algorithm, the value added by this multi-agent formulation comes from two features that were absent in the single-agent implementation: data compartmentalization and information sharing. Data compartmentalization arises from the fact that conformations are labeled and utilized by the agent who discovers them, so each agent learns the parameters to compute the rewards from different data points. This is done with the intention to constrain what information will be used to score the conformations in a region of the landscape. Moreover, the agents share information at the action-space discretization step to signal to others what conformations should be deprioritized based on the structures they have already observed.

The idea of utilizing prior known states of a system to accelerate MD simulations has been explored in the past. For instance, FAST and TSMD use scoring functions that can be set to the RMSD between the current conformation and the desired final state to guide simulations. Nonetheless, the goal of Multi-agent REAP is not to observe transitions between predetermined conformations; if two agents start exploring a landscape from different initial states, their exploration efficiency may improve by interacting with each other to avoid resampling observed regions, but no assumptions are made about the new states that may be discovered.

Tangent Space Least Adaptive Clustering (TSLC) is an algorithm tightly related to REAP but whose clustering and reward schemes are better suited to handle nonlinear CVs.^37^ Given that this algorithm belongs to the family of adaptive sampling regimes that perform a discretization of the action space, our multi-agent formulation can be easily extended to this algorithm. To show the flexibility of the main idea behind our multi-agent formulation, we also introduce a multi-agent version of TSLC (Multi-agent TSLC) and compare it against its single-agent baseline.

The rest of this paper is organized as follows: the method section describes the modified versions of REAP and TSLC. Then, performance comparisons are made on artificial toy potentials to illustrate the behavior of the multi-agent algorithms. Following, a comparison between Multi-agent REAP and AdaptiveBandit is done using MD simulations of alanine dipeptide. Lastly, we compare the original and multi-agent REAP algorithms on realistic systems (Src kinase,^4^ OsSWEET2b^38^) using kinetic Monte Carlo (KMC) sampling based on data from previous studies. We found that in all cases the multi-agent algorithms were able to more quickly explore the free energy landscape compared to their single-agent counterparts and AdaptiveBandit.

## 2 Methods

### 2.1 Multi-agent REAP algorithm

Algorithm 1 shows the outline for Multi-agent REAP. Since this method shares most of its logic with the original REAP algorithm,^28^ we will focus on describing the changes in the multi-agent formulation.

#### Algorithm 1: Multi-agent REAP

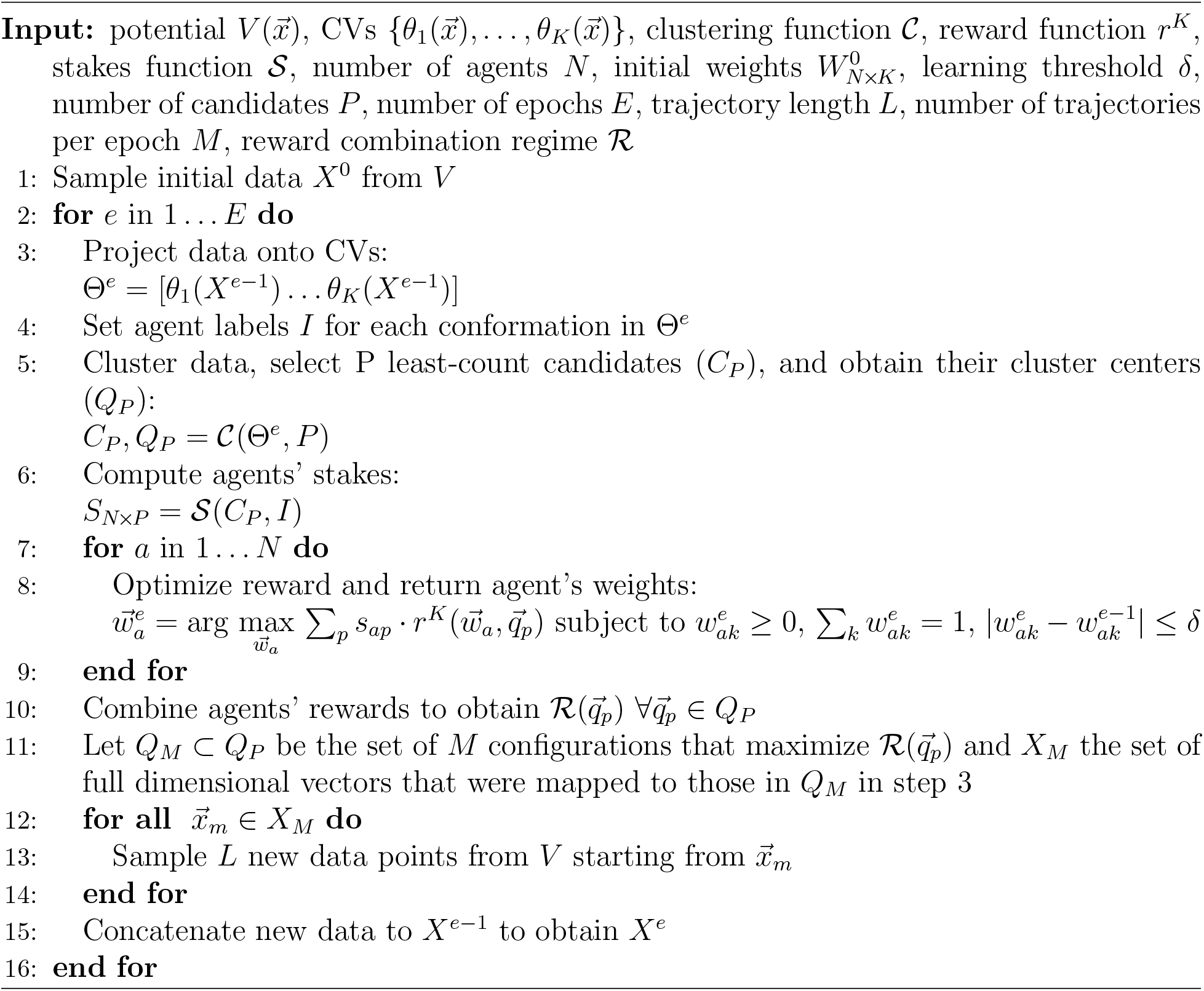

The first notion to discuss is the definition of an agent. In general, an agent is an object that can collect observations from an environment. Based on its observations, the agent falls into a specific state from a potentially infinite state set. The policy function will then map the agent’s state into the next set of actions. The environment will return new observations dependent on the actions taken. To guide the behavior of the agent, a reward function is defined such that it dictates the development of the agent’s policy.^25^ Namely, the policy will select actions that optimize the current or future rewards.

For Multi-agent REAP, the definitions of environment, state, and action remain unchanged in comparison to the original REAP algorithm. The environment is usually the Hamiltonian of a many-body system. An important feature of such a Hamiltonian is that its free energy landscape lies near a low-dimensional manifold. This low-dimensional space is spanned by a set of CVs. Each REAP agent can rank user-given CVs to prioritize a direction of exploration, but no new variables can be discovered. The agents can make observations by sampling a thermodynamic ensemble of the system using standard MD methods. The state of an agent is defined as the set of conformations that the agent has discovered. An action is defined as a conformation from which the agent can launch an MD trajectory.

The reward function of Multi-agent REAP differs from that of the original implementation. To explain the difference between the single- and multi-agent reward functions we introduce the concept of an agent’s *stake* in an action. REAP relies on a clustering step where the action space is reduced and discretized. In other words, the large amount of possible conformations from which to launch new simulations is reduced to a relatively small number of representative clusters. Usually, the centroids of the clusters conform the reduced action space. In Multi-agent REAP, the data from all agents is pooled together during action-space discretization (step 5 in algorithm 1), so agents are effectively sharing information. Moreover, each agent possesses a “stake” in each cluster. The stake 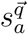 that agent *a* has on action 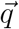 is computed as a function of the number of conformations discovered by agent *a* classified into the cluster corresponding to action 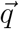. A simple example of a stake function is one that returns the fraction of frames in the cluster of action 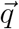 that belong to agent *a*. We propose other alternatives (see equations 3–6), but note that the choice among these functions does not dramatically alter the results (see Supporting results). Stakes are intended to fulfill the condition 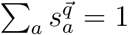 for any action 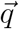. In essence, Multi-agent REAP uses each agent’s stake to weight the reward that it feels from a given cluster. Since the REAP algorithm is intended to encourage exploration, the reward is proportional to the standardized Euclidean distance between the action and the mean of the discovered conformations in CV space. Although the use of different distance functions is possible, this metric is retained by our multi-agent formulation.

The single-agent reward function is defined as

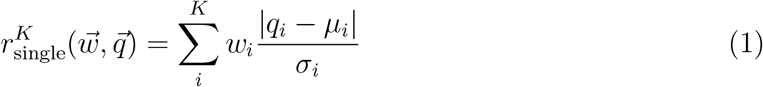

where *K* is the number of CVs, *w_i_* is the weight assigned to CV *i*, 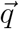 is a cluster center (in CV space), 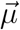 is the mean of all the discovered conformations, and 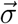 is the standard deviation. Building from equation 1, the reward an agent assigns to an action is simply expressed as

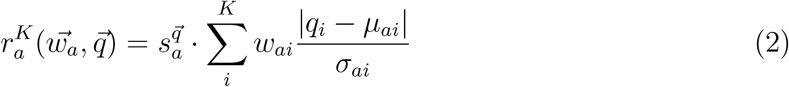

where 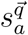 is the stake that agent *a* has on cluster center 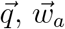 are agent’s *a* weights, and 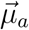 and 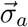 are the mean and standard deviation of all the conformations discovered by agent *a*. The stake is computed using a function 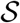. The proposed forms of 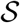 are

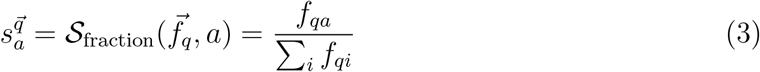

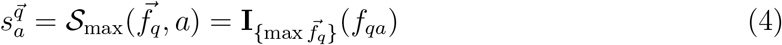

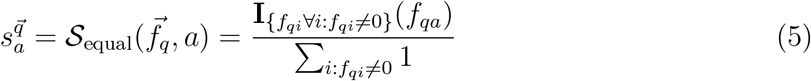

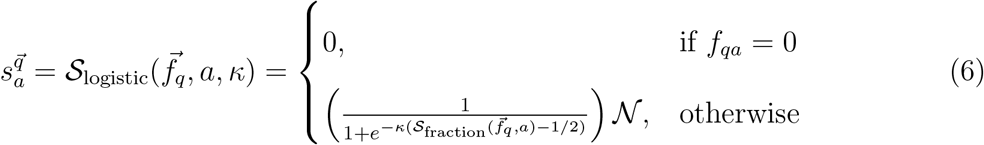

where the length of 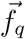 is the number of agents and each entry contains the number of frames classified as action 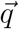 that each agent discovered, **I** is the indicator function, *κ* is a tunable parameter, and 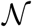 is a normalization factor. In simple terms, 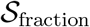 returns the fraction of conformations that agent *a* possesses in action 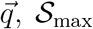 assigns a stake of 1 to the agent with the largest number of conformations in the action (and 0 to the rest), 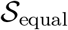 assigns equal stakes to all agents with at least one frame in the cluster, and 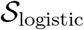 provides the option of a tunable function that can produce intermediate behaviors according to the value of *κ* (for large *κ* it approximates 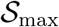 and for small *κ* it approximates 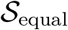).

Once the agents’ stakes have been assigned, the CV weights are updated by optimizing the reward. It is important to highlight that the agents only share information during action-space discretization. For this reason, all other parameters remain constant when updating the weights. Since the reward function only depends on the weights from the given agent, we can carry out agent-wise optimization at step 8 of algorithm 1. 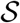 affects the reward optimization by determining how much weight each cluster will carry according to the relative representation of each agent. Employing 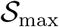 would result in agents only optimizing the reward with respect to the clusters where they are highly represented, whereas utilizing 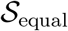 would allow the agents to equally weight all the clusters they have observed at least once. 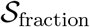 results in the clusters being weighted in direct proportion to the representation of the agent in the cluster. Results of 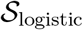 will depend on the value of *κ*. For a comparison among these functions see the Supporting results. 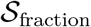 was used in all the tests unless otherwise noted.

In regards to the policy followed to choose the actions, the single-agent implementation simply selects the top *M* actions when ranked by their reward. However, the multi-agent implementation must deal with the nontrivial question about how to combine the rewards from different agents. We propose three ways in which the actions can be selected. The proposed schemes to combine the rewards are

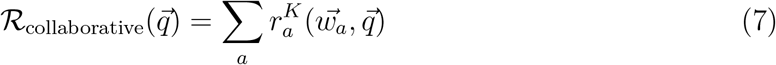

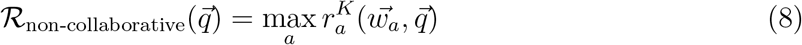

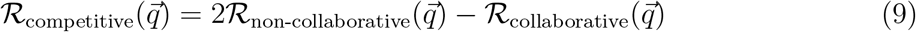

In equation 7, the rewards from all agents are added together. In the RL literature, such schemes where rewards are added to select the most optimal global actions are termed *collaborative* regimes.^39^ On the other hand, equation 8 sets the reward of the action as the maximum reward assigned by a single agent. We term this regime *non-collaborative*. Finally, we proposed a *competitive* regime where the global reward of an action is the maximum reward set by an agent minus the reward assigned by all others. Intuitively, the exploration of a free energy landscape can be framed as a collaborative task; if an action is deemed to be highly rewarding by all agents, then it should be selected. We cannot discard the (albeit counterintuitive) possibility that a competitive regime may lead to better performance in settings where actions that are deemed rewarding by multiple agents should be discouraged. Nonetheless, we did not detect a significant difference between reward combination regimes in our tests (see Results and discussion).

The new data generated after selecting action 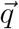 is assigned to the agent that possesses the highest stake in that action. In other words, the agent whose stake is maximal for a selected action is the one that “executes” it. A consequence of this decision is that the agents that are the most “proficient” at exploring the landscape will continue receiving higher rewards and executing new actions. Therefore, computational resources are distributed unequally across agents.

### 2.2 Multi-agent TSLC algorithm

The multi-agent REAP algorithm shows improved performance in systems with linear and orthogonal CVs (see Results and discussion). However, when the relevant CVs that describe the slow dynamics of the system are not linear, a different method to derive the weights may be better suited. Tangent Space Least Adaptive Clustering (TSLC) was introduced as an extension of REAP precisely to handle these cases.^37^ However, despite its ability to capture non-linear CVs, TSLC shows limitations similar to those of single-agent REAP, and for this reason we introduce Algorithm 2. This extension of TSLC is analogous to that presented in Algorithm 1 for REAP.

TSLC utilizes a distinct clustering function termed Clust.^37^ The clustering of the data is independent of which agent discovered each conformation (i.e., the data is shared among agents during this step as in Multi-agent REAP). The distribution parameters that the agents use to obtain their rewards are replaced by a matrix *A_a_* that is computed in the same way as in Buenfil *et al.*^37^ except for the fact that the stake modulates how the cluster is weighted by the agent (step 10 in Algorithm 2). The matrix *A_a_* is constructed by first approximating the local tangent space of each cluster (using PCA, step 7), finding the gradients of the CVs at each cluster center (step 8), and then adding the contribution of each cluster in step 10. In step 11, we find the linear combination of the gradients of the CVs that maximize the projection onto the tangent spaces. This maximization is expressed as an eigenvalue problem and the final weights are the squared entries of the eigenvector associated with the largest eigenvalue. The last steps are identical to those in Algorithm 1, noting that the values 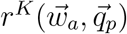 are computed to select the actions but not to optimize the CV weights.

#### Algorithm 2: Multi-agent TSLC

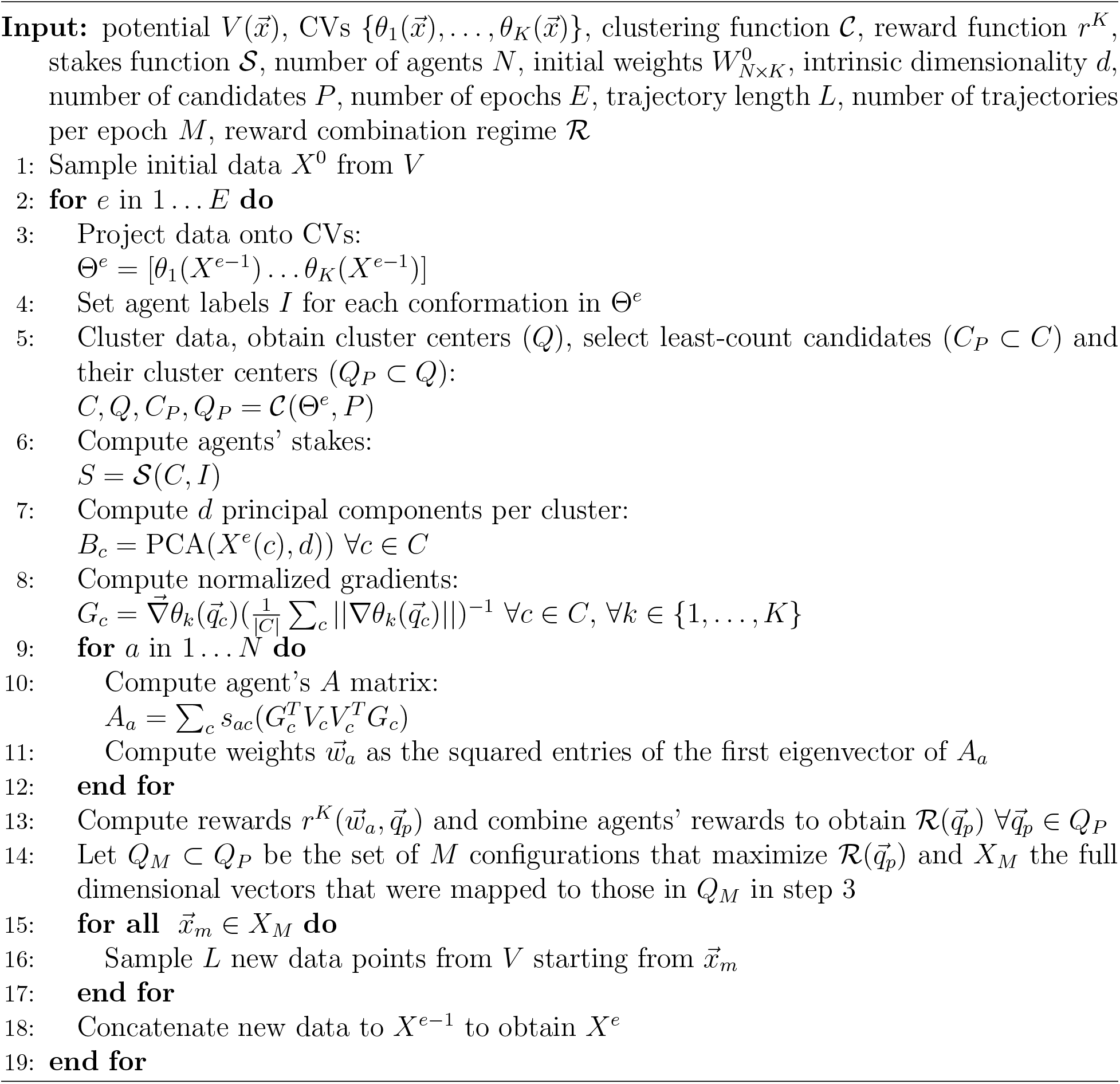

## 3 Results and discussion

### 3.1 Cross potentials

We begin by showing the behavior of Multi-agent REAP in two idealized potentials where a single particle diffuses following a Langevin dynamic.^28^ These examples represent “adversarial” landscapes for Single agent REAP because they showcase its shortcomings in comparison to the multi-agent implementation. LC runs were also performed and are used as baselines to compare the algorithms. Details of the simulations are described in Supporting methods.

Figure 2 shows the results of comparing the algorithms in a *symmetric* cross potential. The potential is shaped like a cross; diffusion along the *x* variable is more relevant for the exploration of the horizontal arms, while the *y* variable is more relevant for the exploration of the vertical arms. The agents are initially placed on the horizontal arms (see Figure 2a). Figure 2c shows the difference in area explored (defined as difference in normalized number of grid points discovered) between Multi-agent REAP and LC. The difference between Single agent REAP and LC is also plotted. The error bars show the 95% confidence intervals after 500 repetitions of the simulations. There are statistically significant differences among all methods; while Multi-agent REAP is able to explore a larger area compared to LC, the single-agent implementation performs worse after roughly 10 epochs. The curves show that around epoch 10, Multi-agent REAP reaches the maximum difference in area explored with respect to LC (approximately 10%) and after that both methods start to converge (as expected for long enough simulation times). However, the Single agent REAP implementation continuously selects actions at the edge of the horizontal arms (Figure 2b), which results on the lack of exploration of the vertical arms. On the other hand, the snapshot of the Multi-agent REAP run shows that the agent starting on the left arm reached the extremes of the vertical arms (and therefore, the other agent does not have the need to launch trajectories in those areas).

**Figure 2:**
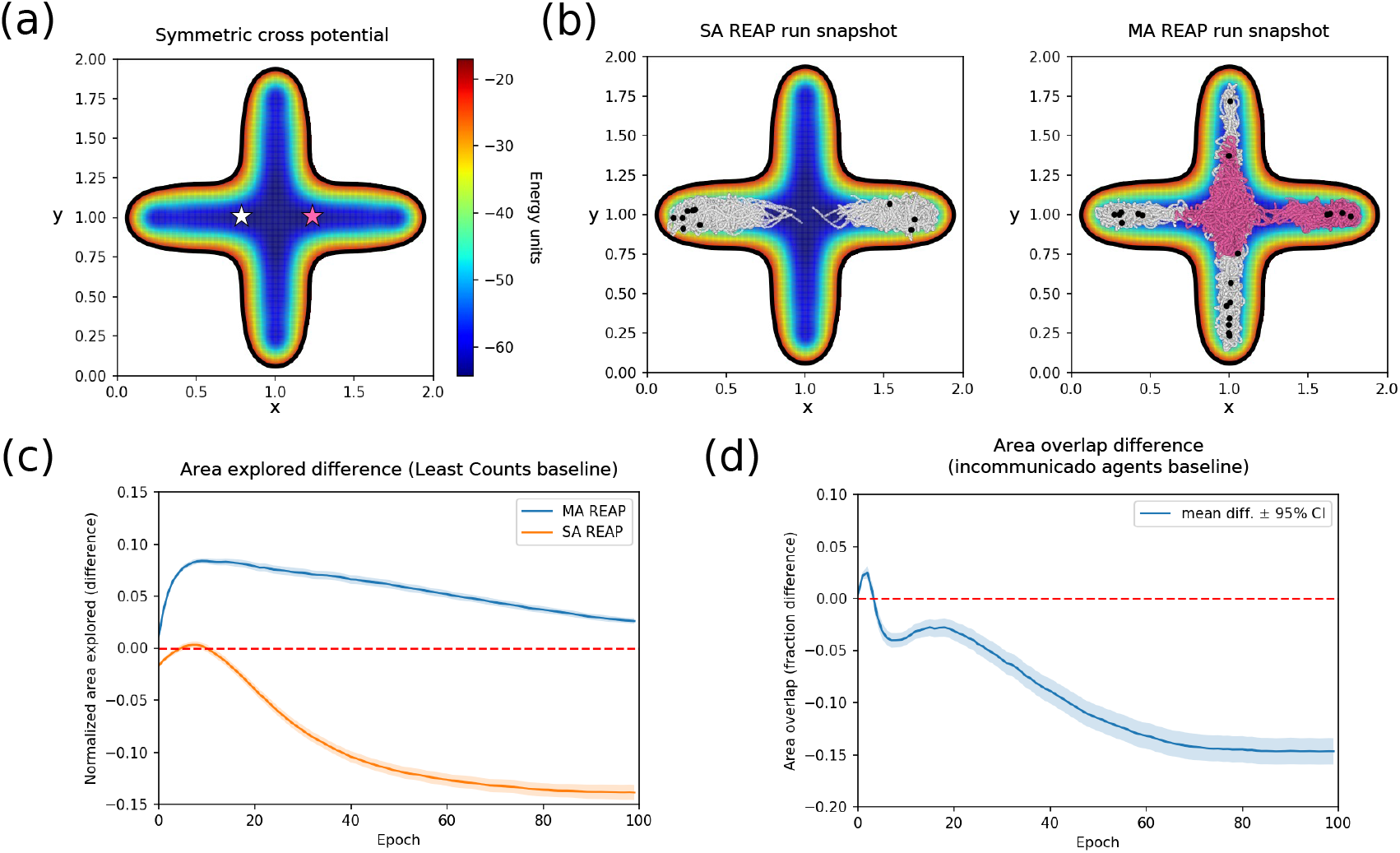
Results on the symmetric cross potential. (a) Plot of the free energy landscape. The white and pink stars mark the initial positions of the agents (both positions are observed by the unique agent in Single agent REAP). (b) Snapshots of the trajectories generated by Single agent REAP (left) and Multi-agent REAP (right) after a full run. Trajectories are colored according to the agent who executed the simulation. Black dots indicate the last set of selected actions. (c) Comparison of normalized area explored. The blue and orange curves show the differences (mean ±95% confidence interval) between the areas explored by Multi-agent REAP or Single agent REAP and LC. (d) Comparison in overlap area between agents. Overlap is defined as the fraction of grid points that both agents have observed over the total number of explored grid points. The curve shows the difference (mean ±95% confidence interval) between Multi-agent REAP and two incommunicado agents.

One may argue that a user of the original REAP algorithm would simply start two independent REAP runs that do not share any information at each starting conformation (i.e., *incommunicado* agents), so that exploration would not become hampered by the greedy selection of actions at the extremes of the *x* variable. But in this case, the agents redundantly explore the same areas of the landscape (see Figure 2d), thus wasting computational resources.

Employing different reward combination regimes did not yield differences (see Figure S2).

Figure 3 shows the comparison results on an *asymmetric* cross potential. This potential is similar to the symmetric one but the minimum at the left extreme of the horizontal arm is deeper than the one on the right. The initial positions of the agents is set to the extremes of the horizontal arms. In this comparison, it is noticeable once again that the Multi-agent REAP algorithm is more efficient at exploring the landscape than the LC adaptive sampling scheme (Figure 3c), while the Single agent REAP implementation suffers from similar non-optimal behavior as in the previous example. However, in this case, the reason for this difference in performance stems from the ability of Multi-agent REAP to allocate more actions to agents that discover new states and earn higher rewards. As the snapshots in Figure 3b show, the agent starting at the right arm is better-poised to explore the landscape in comparison to the left agent, which starts in a deep free energy minimum (e.g., a kinetic trap or absorbing state). Given that the agent starting on the right explores a wider area, it possesses all the stakes in the newly discovered sates, it receives higher rewards, and it is assigned more actions as the run continues (Figure 3d).

**Figure 3:**
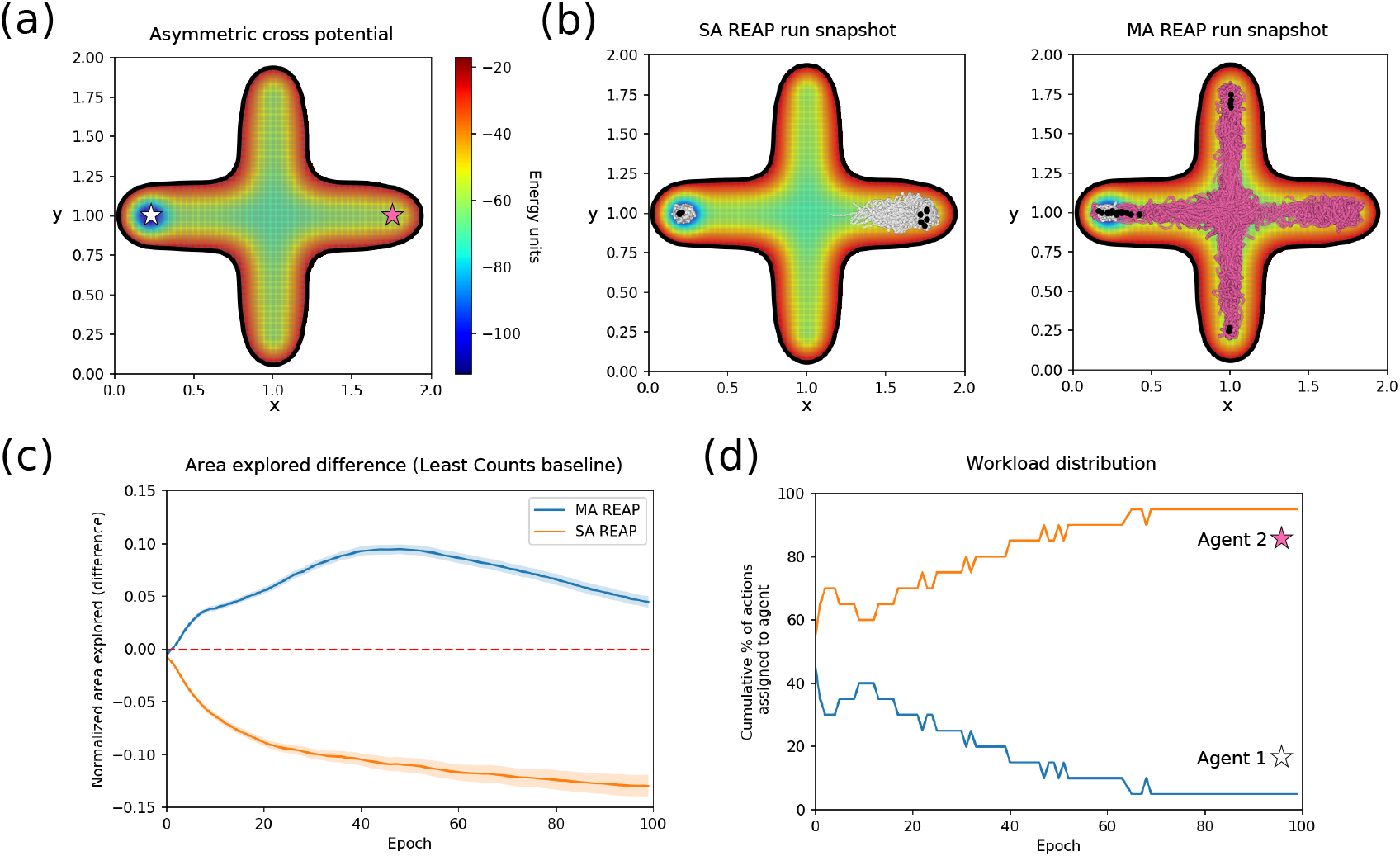
Results on the asymmetric cross potential. (a) Plot of the free energy landscape. The white and pink stars mark the initial positions of the agents (both positions are observed by the unique agent in Single agent REAP). (b) Snapshots of the trajectories generated by Single agent REAP (left) and Multi-agent REAP (right) after a full run. Trajectories are colored according to the agent who executed the simulation. Black dots indicate the last set of selected actions. In the multi-agent snapshot, the agent that starts at the right arm explores most of the landscape because it is assigned more actions. (c) Comparison of normalized area explored. The blue and orange curves show the differences (mean ±95% confidence interval) between the areas explored by Multi-agent REAP or Single agent REAP and LC. (d) Cumulative percentage of actions assigned to each agent after the given epoch for the run plotted in part (b).

In summary, our multi-agent implementation retains the ability of REAP to sample landscapes where there is a clear advantage to preferentially sample along a relevant variable. This is demonstrated by the better higher exploration area of Multi-agent REAP when compared to LC adaptive sampling. Moreover, our algorithm addresses the shortcomings that Single agent REAP presented when attempting to utilize information from distinct regions of the landscape simultaneously. This is evidenced by the ability of Multi-agent REAP to compartmentalize information (the agents only learn from data that comes from their explored region), reduce redundant sampling (area overlap between agents is diminished in comparison to independent agents), and distribute the workload unequally (the agent that explores more is assigned more actions).

### 3.2 Alanine dipeptide

In this section, we simulate alanine dipeptide, a usual example of an all-atom system typically employed to compare MD simulation methods. We compare the performance of Multi-agent REAP against continuous MD and another adaptive sampling regime based on the classical multi-armed bandit algorithm (AdaptiveBandit).^30^ For details on the simulations, see the Supporting methods.

Figure 4 shows the results of the comparison among the three simulation methods. The CVs used for exploration were the *ϕ* and *ψ* dihedral angles (for both Multi-agent REAP and AdaptiveBandit). We observed that, for a total simulation time of 16 ns, Multi-agent REAP was able to reach and thoroughly explore the state with *ϕ* > 0 (Figure 4, bottom panels). This state was not reached by a continuous MD simulation of the same length (Figure 4, top panels). Although AdaptiveBandit sampled a low number of frames in this state, the algorithm did not readily select such structures to restart simulations (Figure 4 middle panels). The result was a largely unexplored area of the landscape.

**Figure 4:**
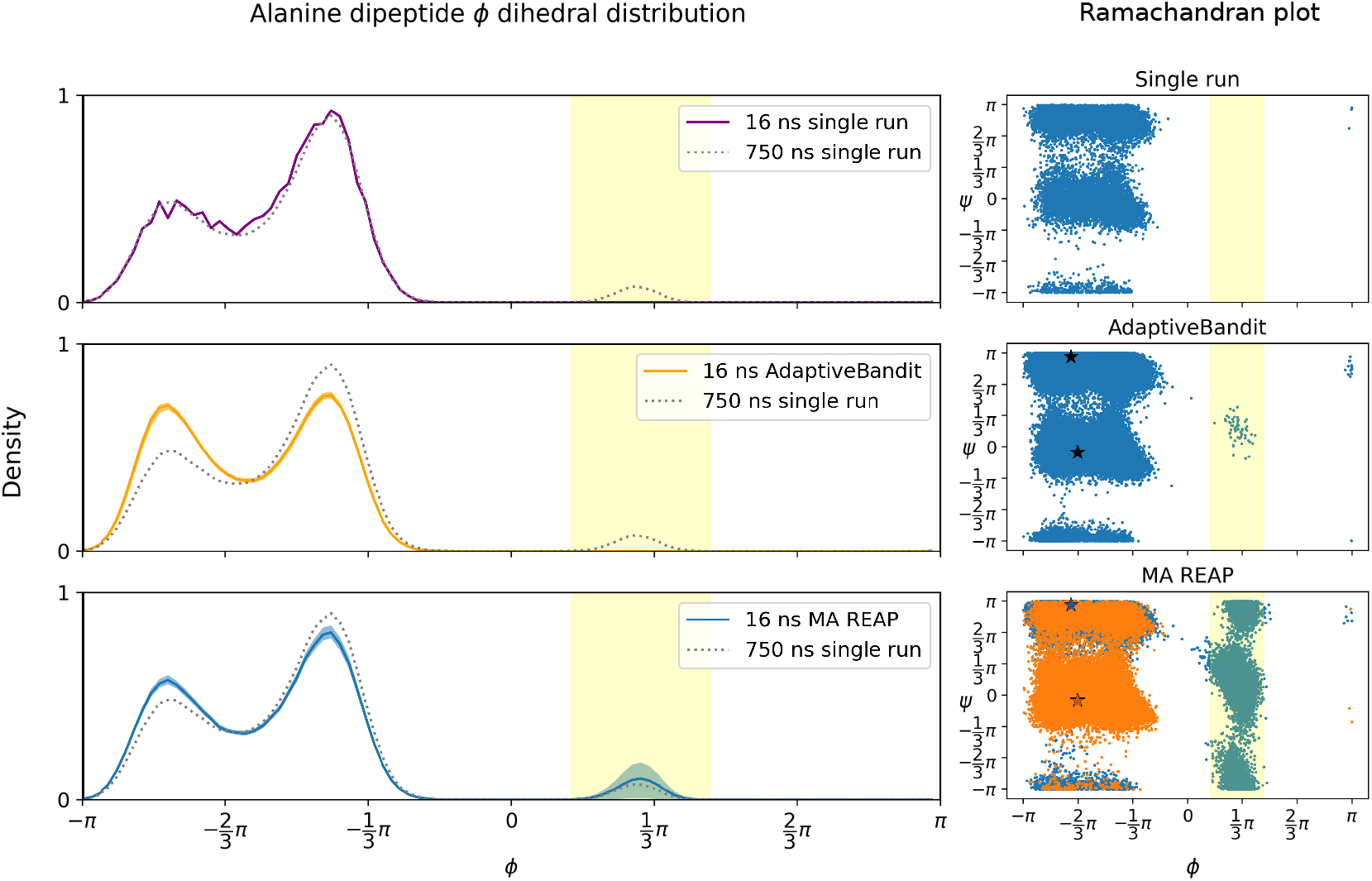
Alanine dipeptide simulation results. (Left panels) Probability density distribution along the *ϕ* dihedral angle for a single run (top), AdaptiveBandit (middle), and Multi-agent REAP (bottom). Curves show mean ± standard error across replicates. (Right panels) Snapshots after a total of 16 ns of MD simulation. The stars represent the starting configurations for the Multi-agent REAP agents and the AdaptiveBandit run. For Multi-agent REAP (bottom), the frames obtained by different agents are colored differently. Multi-agent REAP thoroughly explores the highlighted region, while a continuous MD and AdaptiveBandit fail to sample this area.

The result is likely due to the fact that the reward function for AdaptiveBandit balances the exploration of new regions of the landscape with the exploitation of known metastable states, as reflected by its binomial reward function.^30^ In this case, the presence of highly stable states with *ϕ* < 0 favors the exploitation of the well-characterized actions over the exploration of the states with uncertain rewards. Since AdaptiveBandit is grounded on the basis of the upper confidence bound (UCB1) algorithm, this sampling scheme presumably obeys the optimal theoretical bound on its regret *L_t_*, 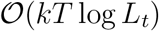.^30^ However, optimally bounding this definition of regret is not necessarily the reason an adaptive sampling technique is utilized. Given that the reward function from AdaptiveBandit is defined as the expected value of the negative free energy, actions that do not achieve the absolute free energy minimum (i.e., the optimal action) will be assigned lower action-values. Nonetheless, MD practitioners often desire to enhance the sampling of rare transitions, excited states, or off-equilibrium processes which are associated with high energies. Therefore, it is favorable for a wide variety of applications to sacrifice the theoretical upper boundary on the regret to accelerate the sampling of such high-energy states.

A user of Multi-agent REAP would hypothetically harness the ability of our algorithm to quickly explore the landscape of the system to later execute an “exploitation” step, where longer, continuous simulations are launched from the observed states to improve the sampling statistics. Similar workflows have been employed in studies such as Zimmerman *et al.*^2^

We can also observe that the probability density distribution along the *ϕ* CV in both AdaptiveBandit and Multi-agent REAP deviates from the equilibrium value (computed from three 250 ns continuous MD simulations retrieved from mdshare^40,41^). Nonetheless, after employing an adaptive sampling scheme, the probability density of an observable would be reweighted through a Markov state model (MSM) to eliminate the statistical bias caused by the repeated restart of simulations from selected states.^42^

### 3.3 Src kinase

In this section, we compare the performance of Single agent REAP and Multi-agent REAP employing Src kinase, a realistic system involved in critical signalling pathways whose malfunction is associated with cancer.^43^ We utilize a previously constructed MSM^4^ to carry out KMC simulations. For details about the simulations, see Supporting methods. The discovery of intermediate states in this system is particularly relevant because they may exhibit allosteric sites that can be targeted by drug design but that are absent in the active or inactive conformations.^4^

Figure 5 shows the results of the KMC simulations. Simulations were started from the active and inactive conformations of the kinase (Figure 5a,b), mimicking the initial information that a researcher would possess from resolved crystal structures.^44,45^ The two CVs that are utilized to project the MSM states are the RMSD of the A-loop and the distance between K295 and E310.^4^

**Figure 5:**
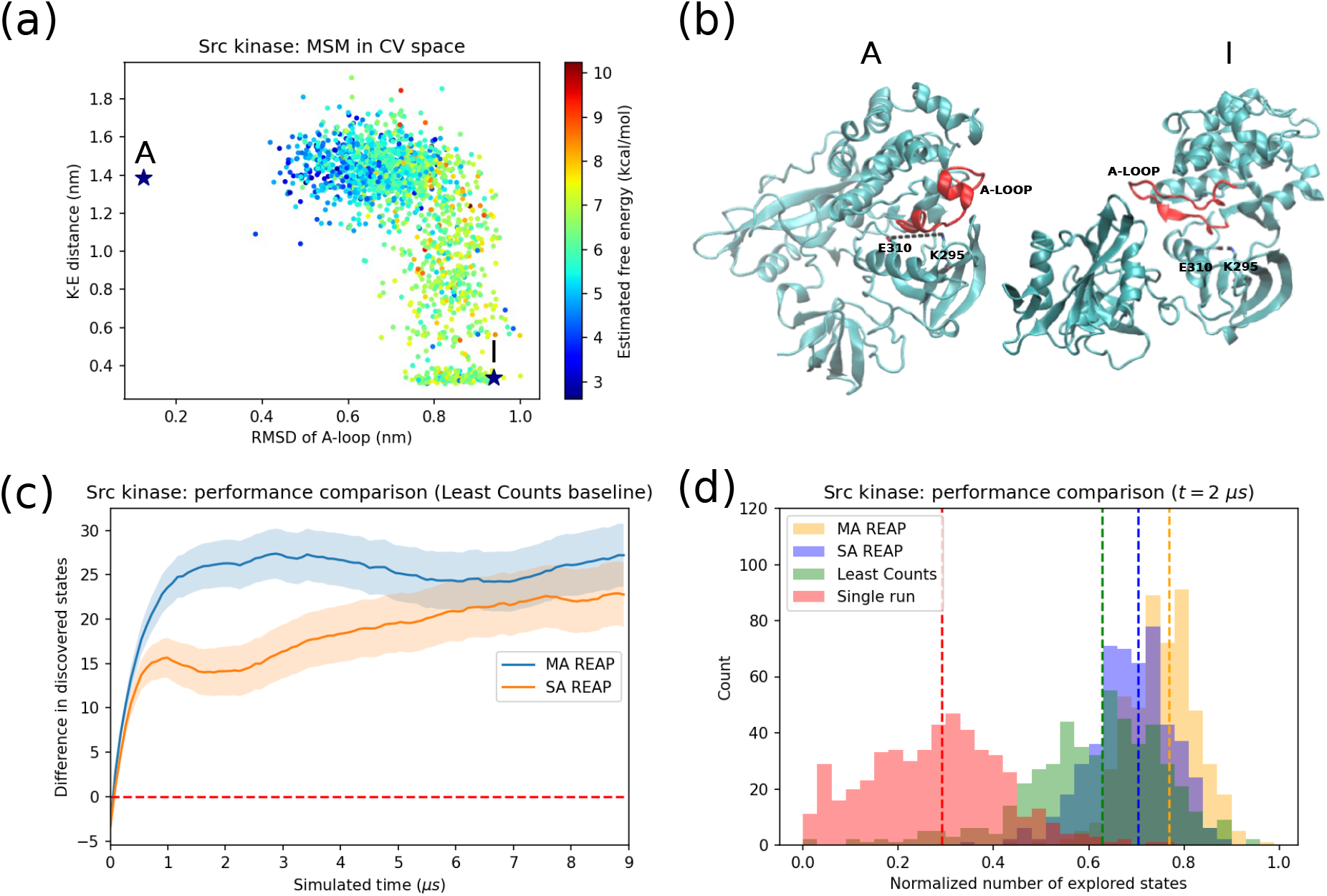
KMC simulation results on Src kinase. (a) Projection of the MSM states on two collective variables (*x*-axis: RMSD of A-loop, *y*-axis: distance between K295 and E310). The stars represent the starting conformations for the simulation. (b) Representative structures of the active (A) and inactive (I) states. The A-loop is colored in red. K295 and E310 are labelled. (c) Performance difference between adaptive sampling schemes. The *y*-axis corresponds to the difference in number of states discovered between MA or Single agent REAP and LC. The plotted curve is the mean ±95% confidence interval for 500 repeats. (d) Histogram of the normalized number of states discovered by single MD simulation runs, LC, Single agent REAP, and Multi-agent REAP after 2 *μ*s of simulation. Normalization is with respect to the maximum number of discovered states.

Figure 5c shows the difference in the number of states discovered between Multi-agent REAP (blue curve) or Single agent REAP (orange curve) and LC adaptive sampling. Both SA and Multi-agent REAP discover more intermediate states than LC for the same simulation time, but Multi-agent REAP outperforms Single agent REAP until both methods converge around 6 *μ*s (all methods should converge for long enough simulation times).

Figure 5d shows the distribution of number of states explored (normalized to the maximum achieved) for the 500 trials after 2 *μ*s. The vertical dashed lines show the means of the distributions. We can observe that Multi-agent REAP achieves a higher average performance than Single agent REAP and LC. Unsurprisingly, both single- and multi-agent REAP improve upon the continuous MD run performance.

Unlike the results in Figure 2c and Figure 3c, Single agent REAP performs better than LC (Figure 5c,d). There are two reasons for this. Firstly, the cross potentials were specifically crafted as adversarial examples for Single agent REAP, so it is expected that the single-agent algorithm will perform particularly poorly. Secondly, KMC simulations allow the system to change states discretely rather than simulating Langevin diffusion. Therefore, selecting actions that present extreme values of the CVs may still result in transitions that uncover intermediate states, avoiding continuously launching unproductive trajectories.

We can also observe that it takes roughly 1 *μ*s for Multi-agent REAP to display its full advantage with respect to other methods (Figure 5c). This is likely due to the time it takes agent 2 (initially in the inactive conformation) to learn advantageous weights (Figure S5). For this agent, moving along the K-E distance CV leads to the discovery of more intermediate states.

In summary, we have shown that Multi-agent REAP discovers intermediate states more efficiently compared to REAP and LC in a KMC simulation of a realistic system.

### 3.4 OsSWEET2b

OsSWEET2b is a vacuolar glucose transporter in rice.^46,47^ The study of this system is relevant to the improvement of crop yields.^48^ Similarly to our Src kinase comparison, we perform KMC on an MSM previously built on the glucose-bound transporter.^38^ Unlike the previous example, the landscape does not present the typical L-shape where REAP is expected to carry a clear advantage.^28^ This is due to the existence of hourglass-like intermediate states.^38,49,50^

We start our simulations from the inward facing (IF) and outward facing (OF) states (Figure 6a,b). Details of the simulations are described in Supporting methods.

**Figure 6:**
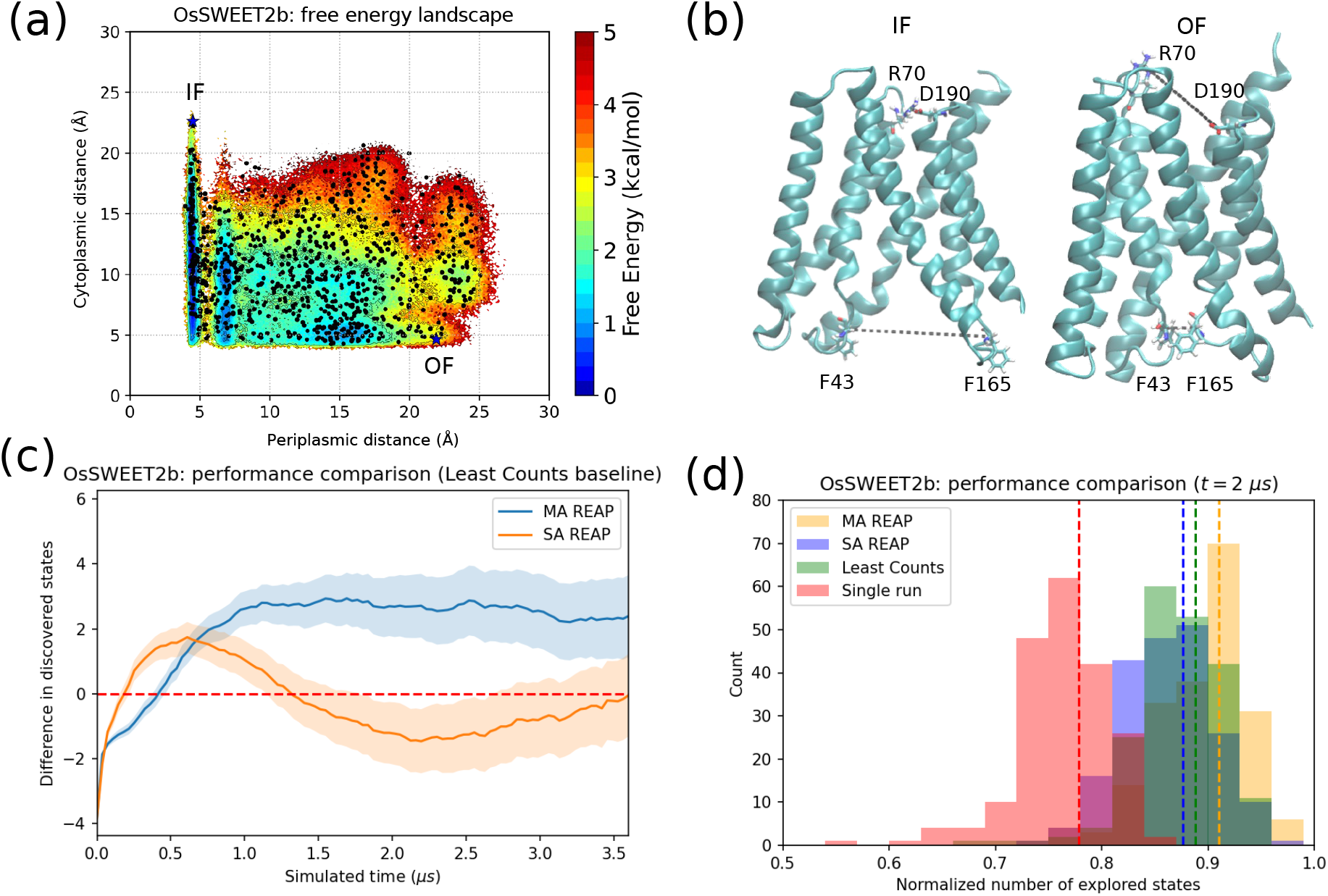
KMC simulation results on OsSWEET2b Markov state model (MSM). (a) Projection of the MSM states on two collective variables: *x*-axis: periplasmic distance (distance between residues D190 and R70), *y*-axis: cytoplasmic distance (distance between F165 and F43). The stars represent the starting conformations for the simulation. (b) Representative structures for inward-facing (IF) and outward-facing (OF) states. (c) Performance difference between adaptive sampling schemes. The *y*-axis corresponds to the difference in number of states discovered between MA or Single agent REAP and LC. The plotted curve is the mean ±95% confidence interval for 200 repeats. (d) Histogram of the normalized number of states discovered by single MD simulation runs, LC, Single agent REAP, and Multi-agent REAP after 2 *μ*s of simulation. Normalization is with respect to the maximum number of discovered states. The *x*-axis starts at 0.5 for clarity.

As a consequence of the characteristics of this system, the advantage of REAP and Multi-agent REAP compared to single runs is less accentuated (Figure 6d). Nonetheless, both SA and Multi-agent REAP still show better performance than continuous MD (defined as number of states discovered). Single agent REAP performs worse or no better than LC after approximately 1.5 *μ*s, while Multi-agent REAP maintains a modest advantage (Figure 6c).

Although Multi-agent REAP shows generally better performance than LC and its single-agent counterpart, it shows a small disadvantage at *t* < 0.5 *μ*s. Such initial setback can be attributed to the time that it takes the agents to learn advantageous weights to facilitate exploration (see Figure S6b). Namely, agent 2 (which starts at the outward facing state) only starts prioritizing state transitions along the cytoplasmic distance CV after roughly 0.5 *μ*s. Before that, the agent observes mostly transitions along the periplasmic distance CV. Conversely, agent 1 (initially at the inward facing state), only begins prioritizing exploration along the periplasmic distance CV after roughly 1 *μ*s. Before that time, this agent observes mostly transitions involving changes in the cytoplasmic distance CV. On the other hand, the weights for the single agent (Figure S6a) show a fluctuating behavior until *t* =1.5 *μ*s, after which the periplasmic distance CV is prioritized. Rather than improving the performance of the agent, these weights harm its ability to explore new states as shown by the drop in performance after 1.5 *μ*s. This is likely due to the over-selection of actions at the extremes of the periplasmic distance variable, which results in unproductive trajectories in a similar way to what was shown in Figure 2 and Figure 3.

In this section, we have shown that Multi-agent REAP is more favorable compared to LC, even in energy landscapes where we would not expect our algorithm to perform better. Moreover, the multi-agent implementation explored more states than the original, single-agent one.

### 3.5 Toroidal potential

In this section, we show that the main idea behind the Multi-agent REAP algorithm can be applied to other adaptive sampling schemes. More specifically, we show that a multi-agent version of TSLC explores a model potential more efficiently than the original algorithm when we allow for multiple starting points in a Langevin diffusion simulation. For details about the simulation, see the Supporting methods.

Unlike the idealized potentials from Figure 2a and Figure 3a, the potential in Figure 7a, termed toroidal or circular^37^ potential, possesses a slow variable that cannot be expressed as a linear combination of the particle coordinates. The appropriate CV is the angle, *θ* = arctan (*y*/*x*) (with its quadrant-dependent sign), but the weight derivation method employed in algorithm 1 cannot consistently assign the highest weight to this variable.^37^ However, the weight derivation employed in algorithm 2 is better suited to learn the weights for this system thanks to its ability to capture global geometric information from locally estimated tangent spaces.

**Figure 7:**
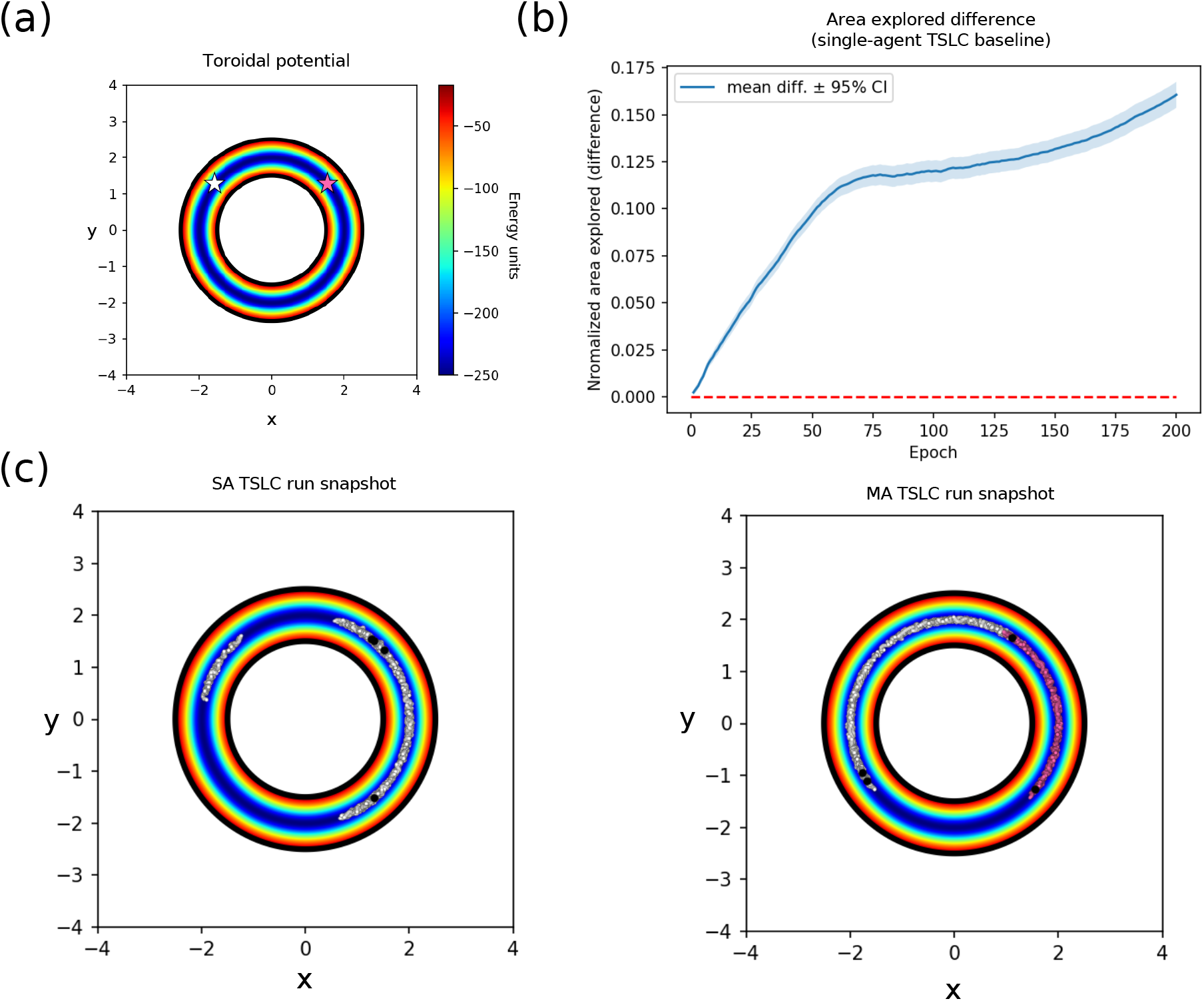
Results on the toroidal potential. (a) Plot of the free energy landscape. The white and pink stars mark the initial positions of the agents (both positions are observed by the unique agent in Single agent REAP). (b) Comparison of normalized area explored. The curve shows the difference (mean ±95% confidence interval) between the areas explored by Multi-agent TSLC and Single agent TSLC. (c) Snapshots of the trajectories generated by Single agent TSLC (left) and Multi-agent TSLC (right) after a full run. Trajectories are colored according to the agent who executed the trajectory. Black dots indicate the last set of selected actions.

Figure 7 shows the result of comparing the original TSLC algorithm to the multi-agent implementation when sampling is started from two initial points, *θ*_1_ = 3/4*π* and *θ*_2_ = 1/4*π* (white and pink stars in Figure 7a respectively). After the single-agent has explored enough of the circle to reach the supplementary angle to either *θ*_1_ or *θ*_2_, TSLC falls into an unproductive loop where actions that are either diametrically opposite in the circle or maximally distant along the *x*-or *y*-coordinate keep getting selected, even though they do not lead to further exploration along the angle CV (Figure 7c, left). On the other hand, Multi-agent TSLC continues selecting actions along the unexplored edge of the circle (Figure 7c, right). Due to this difference in behavior, we observe a clear difference in performance between the two algorithms (Figure 7b). The difference continues to increase as the number of epochs grows because the single-agent implementation does not continue exploring the circle after falling in the unproductive loop. The multi-agent formulation of TSLC preserves the ability to estimate the CV weights like the original algorithm (Figure S7). Employing different reward combination regimes did not yield differences (see Figure S8).

## 4 Conclusion

In this study, we developed adversarial examples where REAP performs worse than LC adaptive sampling and empirically demonstrated that Multi-agent REAP overcomes the limitations of its predecessor. Moreover, we showed that Multi-agent TSLC explores an idealized energy potential more effectively than TSLC. We additionally demonstrate the advantage of Multi-agent REAP in three test systems: alanine dipeptide, an Src kinase, and the transporter OsSWEET2b. We provide evidence to show that Multi-agent REAP more effectively samples the rare state transitions of alanine dipeptide compared to traditional MD simulations or AdaptiveBandit. For Src kinase and OsSWEET2b, we demonstrate that Multi-agent REAP explores more states than LC or Single agent REAP for the same simulation time. Interestingly, for all the cases we tested, the combination regime for the rewards did not affect the results (see Figure S2 and Figure S8). However, there might be landscapes for which the choice of reward combination regime does affect the performance. Optimal selection of 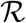 remains a point of investigation.

The number of agents to be used in our algorithms is limited by the number of known distinct states of the system that the researcher can employ as initial coordinates. There are no reasons to expect a better performance from the multi-agent formulations when all agents observe the same initial state. This bears similarity to a limitation present in multi-walker metadynamics, where placing the walkers in the same energy minimum results in suboptimal sampling because the trajectories must become uncorrelated before the landscape converges.^36^

Since we compared intrinsically parallel algorithms, computational power was held constant in the simulations (i.e., the same number of trajectories were launched per epoch regardless of the adaptive sampling method used). The number of trajectories launched in Multi-agent REAP/TSLC is independent of the number of agents *N*, so there is no direct scaling relationship between *N* and the explored area. The benefit of employing the multi-agent algorithms will depend on the prior information on the system (known states) and the nature of its free energy landscape. In this way, the differences in performance that we observed are purely due to algorithmic differences; Multi-agent REAP/TSLC accelerate the exploration due to their ability to compartmentalize and share information, not due to a trivial increase in computational resources.

In comparison to FAST, our methods have the advantage of ranking the CVs according to their relative importance for sampling. Since FAST is not intended to combine information from different CVs, we do not directly compare against this algorithm. A comparison between FAST and AdaptiveBandit is available in the literature.^30^ On the other hand, other unbiased enhanced sampling methods such as PaCS-MD51and TS-MD^29^ are too distant in scope for a comparison to be warranted.

Additionally, we did not compare Multi-agent TSLC with Multi-agent REAP. A comparison between the single-agent versions exists in the literature.^37^ The choice between TSLC or REAP (single- or multi-agent versions) will largely depend on the characteristics of the free energy landscape. TSLC-derived weights will offer an advantage when the CVs are not orthogonal and linear. It is unlikely that this information will be available to a researcher prior to simulating a system. In this regard, the biggest limitation of all CV-based methods is that the important reaction coordinates may be unknown prior to executing the simulations. However, past works explore the possibility of extracting relevant CVs from evolutionary information and applying them in adaptive sampling regimes for proteins.^52^

In terms of future improvements for Multi-agent REAP/TSLC, the reward function for Multi-agent REAP is purely geometrically motivated; rewards are directly proportional to the deviation of conformations from the mean populations. In Multi-agent TSLC, more information about the local topology of the landscape is utilized, but we must make stronger assumptions about the characteristics of the landscape. In both cases, no thermodynamic information is used. In AdaptiveBandit, the free energy is estimated in the reward function to represent the exploitation term, but we observed that this choice hindered exploration (Figure 4). The question of how to encourage exploration has been pondered in the RL literature; perhaps, reformulating the reward function in the form of an entropy-regularized reward^24,53^ might result in a theoretically grounded formulation that guides exploration. Moreover, recent works in multiple-agent coordination^54^ may inspire new formulations of the reward function with improved performance.

In conclusion, we imported the concept of multi-agent RL into the development of adaptive sampling algorithms for MD simulations. We introduced modifications that yielded new algorithms which extend the functionality and improve the performance of REAP and TSLC. More specifically, our multi-agent formulations record which agent discovered each conformation and share this information at the action-discretization step. Moreover, the stakes function modulates how different agents sense the rewards originating from discovered states of the system. These modifications improved exploration performance in all tested cases. Beyond the algorithms proposed here, a key takeaway from this work is that agent coordination can be incorporated into most adaptive sampling strategies, thus opening new avenues to develop methods that are better suited to harness information from different regions of the energy landscape.

## Acknowledgement

D.S. acknowledges support from the National Science Foundation Early CAREER Award (NSF-MCB-1845606). The authors thank James Buenfil, Samson J. Koelle, and Prof. Marina Meila for sharing early-stage TSLC code, which facilitated the implementation of Multi-agent TSLC.

## Supporting Information Available

Supporting Information accompanies this paper online. Code necessary to reproduce the simulations is available on https://github.com/ShuklaGroup/MA_REAP. This material is available free of charge via the Internet at http://pubs.acs.org/.

