## Supporting Information for "Multi-Agent Reinforcement Learning-based Adaptive Sampling for Conformational Sampling of Proteins"

#### 1 Supporting methods

##### 1.1 Langevin dynamics simulations

Langevin dynamics simulations were run using OpenMM 7.7[S1] The Langevin integrator was configured to use a temperature of 300 K, a friction coefficient of 1 $ps^{-1}$, and a time step of 2 fs. The single particle present in the system has a mass of 100 Da.

Three different potential functions were tested with different adaptive sampling schemes (see Main Text).

The symmetric cross potential was defined as:

$$\begin{aligned} V(x,y) = & -25e^{-.5((x-0.2)^2/.008+(y-1)^2/.008)} \\ & -25e^{-.5((x-1)^2/.008+(y-0.2)^2/.008)} \\ & -25e^{-.5((x-1.8)^2/.008+(y-1)^2/.008)} \\ & -25e^{-.5((x-1)^2/.008+(y-1.8)^2/.008)} \\ & -50e^{-.5((x-0.4)^2/.04+(y-1)^2/.02)} \\ & -50e^{-.5((x-0.8)^2/.04+(y-1)^2/.02)} \\ & -50e^{-.5((x-1.2)^2/.04+(y-1)^2/.02)} \\ & -50e^{-.5((x-1.6)^2/.04+(y-1)^2/.02)} \\ & -50e^{-.5((x-1)^2/.02+(y-0.4)^2/.04)} \\ & -50e^{-.5((x-1)^2/.02+(y-0.8)^2/.04)} \\ & -50e^{-.5((x-1)^2/.02+(y-1.2)^2/.04)} \\ & -50e^{-.5((x-1)^2/.02+(y-1.6)^2/.04)} \\ & +60e^{-.5((x-1)^2/.02+(y-1)^2/.02)} \end{aligned}$$

The asymmetric cross potential was defined as:

$$
\begin{aligned}
V(x,y) = &- 80e^{-.5((x-0.2)^2/.008+(y-1)^2/.008)} \\
&- 25e^{-.5((x-1)^2/.008+(y-0.2)^2/.008)} \\
&- 25e^{-.5((x-1.8)^2/.008+(y-1)^2/.008)} \\
&- 25e^{-.5((x-1)^2/.008+(y-1.8)^2/.008)} \\
&- 50e^{-.5((x-0.4)^2/.04+(y-1)^2/.02)} \\
&- 50e^{-.5((x-0.8)^2/.04+(y-1)^2/.02)} \\
&- 50e^{-.5((x-1.2)^2/.04+(y-1)^2/.02)} \\
&- 50e^{-.5((x-1.6)^2/.04+(y-1)^2/.02)} \\
&- 50e^{-.5((x-1)^2/.02+(y-0.4)^2/.04)} \\
&- 50e^{-.5((x-1)^2/.02+(y-0.8)^2/.04)} \\
&- 50e^{-.5((x-1)^2/.02+(y-1.2)^2/.04)} \\
&- 50e^{-.5((x-1)^2/.02+(y-1.6)^2/.04)} \\
&+ 53e^{-.5((x-1)^2/.02+(y-1)^2/.02)}
\end{aligned}
$$

The toroidal potential was defined similarly as in Buenfil *et al.*:[S2]

$$
V(x,y,z) = -250e^{-10\left[\left(\sqrt{x^2+y^2}-2\right)^2+z^2\right]}
$$

For the symmetric and asymmetric cross potential trials, 500 replicates were performed using Least Counts (LC), REAP, and MA REAP. Each run lasted 100 epochs. For REAP and MA REAP, 50 LC candidates were selected per epoch. 20 trajectories were spawn per epoch, each one of a length of 500 time steps (1 ps). The maximum change in CV weights allowed in an epoch was $\delta = 0.02$. For clustering, $k$-means was utilized with random initialization of cluster centers and 5 restarts. $10^4$ frames were randomly sampled from the total accumulated data to perform clustering at each epoch. The number of clusters was set similarly as in Clust[S2] using parameters $b = 3 \times 10^{-4}$, $d = 2$, and $\gamma = 0.6$. Fraction stakes and collaborative rewards were used.

For the toroidal potential trials, 500 replicates were performed using TSLC and MA TSLC. Each run lasted 200 epochs. 50 LC candidates were selected per epoch. 4 trajectories were spawn per epoch, each one of a length of 500 time steps (1 ps). For clustering, Clust[S2] was utilized with parameters $b = 7 \times 10^{-3}$, $d = 4$, and $\gamma = 0.7$. $5 \times 10^5$ frames were randomly sampled from the total accumulated data to perform clustering at each epoch. Fraction stakes and collaborative rewards were used.

#### 1.2 All-atom molecular dynamics simulations

All-atom MD simulations were carried out in OpenMM 7.7.[S1] The Langevin integrator was configured to use a temperature of 300 K, a friction coefficient of 1 $\text{ps}^{-1}$, and a time step of 2 fs. A Monte Carlo barostat was set to maintain the pressure at 1 bar. All bonds involving hydrogen were constrained. AdaptiveBandit runs were performed using ACEMD3[S3] with identical settings.

For alanine dipeptide, the system was sourced from Harrigan *et al.*[S4] The amber99sb force field and the tip3p water model were used to match Shamsi *et al.*[S5] 20 replicates were performed. For each replicate, 200 epochs were run. 4 trajectories of 20 ps were spawn per epoch. For AdaptiveBandit, the default exploration constant $c = 0.01$ was used.[S6] For MA REAP, 12 LC candidates were selected in each epoch. The maximum change in CV weights allowed in an epoch was $\delta = 0.02$. For clustering, $k$-means was utilized with random initialization of cluster centers and 5 restarts. $10^4$ frames were randomly sampled from the total accumulated data to perform clustering at each epoch. The number of clusters was set similarly as in Clust[S2] using parameters $b = 2 \times 10^{-3}$, $d = 2$, and $\gamma = 0.7$. Fraction stakes and collaborative rewards were used.

#### 1.3 Kinetic Monte Carlo simulations

Markov state models (MSMs) for Src kinase and OsSWEET2b were sourced from Shukla *et al.*[S7] and Selvam *et al.*,[S8] respectively. PyEMMA 2.5[S9] was used to generate realizations of the MSMs. This type of simulation is termed kinetic Monte Carlo (KMC) and it consists of sampling state transitions according to the stochastic transition matrix given by the MSM.

For Src kinase, the initial states were selected to correspond to the active and inactive states as described in Shukla *et al.*[S7] 500 replicates were performed. For each replicate, 100 epochs were run. 2 trajectories of 9 steps were spawn per epoch. Since the lag time of the MSM is 5 ns, each trajectory is equivalent to 45 ns of simulated time. 12 LC candidates were selected in each epoch. The maximum change in CV weights allowed in an epoch was $\delta = 0.1$. For clustering, $k$-means was utilized with $k$-means++ initialization of cluster centers and 3 restarts. The number of clusters was set to the number of discovered states. Max stakes and collaborative rewards were used.

For OsSWEET2b, the initial states were selected to correspond to the inward-facing and outward-facing states as described in Selvam *et al.*[S8] 200 replicates were performed. For each replicate, 100 epochs were run. 2 trajectories of 1 step were spawn per epoch. Since the lag time of the MSM is 18 ns, each trajectory is equivalent to 18 ns of simulated time. 12 LC candidates were selected in each epoch. The maximum change in CV weights allowed in an epoch was $\delta = 0.01$. For clustering, $k$-means was utilized with $k$-means++ initialization of cluster centers and 3 restarts. The number of clusters was set to the number of discovered states. Max stakes and collaborative rewards were used.

### 2 Supporting results

#### 2.1 Comparison of stakes functions

In order to compare the performance of different stakes functions (equations 3–6 in Main Text), we ran additional simulations on the symmetric cross potential. Identical settings and parameters as those described in Langevin dynamics simulations were used, with the exception that logistics stakes, $\mathcal{S}_{\text{logistic}}$ (equation 6 in Main Text), with different $\kappa$ values were employed. 100 replicates were run for each $\kappa$ value. This represents a thorough comparison among our proposed stakes functions because for $\kappa = 0.1$, $\mathcal{S}_{\text{logistic}} \approx \mathcal{S}_{\text{equal}}$ and for $\kappa = 100$, $\mathcal{S}_{\text{logistic}} \approx \mathcal{S}_{\max}$ (see Figure S9a).

Figures S9c,d show the differences in performance with respect to the "neutral" choice of fraction stakes (equation 3 in Main Text). From Figure S9c, we can conclude that using different stakes functions does not dramatically alter the explored area. We observed a statistically significant difference for $\kappa = 100$ (in favor of fraction stakes), but the magnitude of the difference is too small to be relevant. On the other hand, we observed that utilizing $\kappa \leq 3$ generally leads to less area overlap between agents in comparison to fraction stakes (Figure S9d). This is indicative that $\mathcal{S}_{\text{equal}}$ or $\mathcal{S}_{\text{logistic}}$ with small $\kappa$ reduce the selection of actions in the same regions for the two agents.

### 3 Supporting Figures

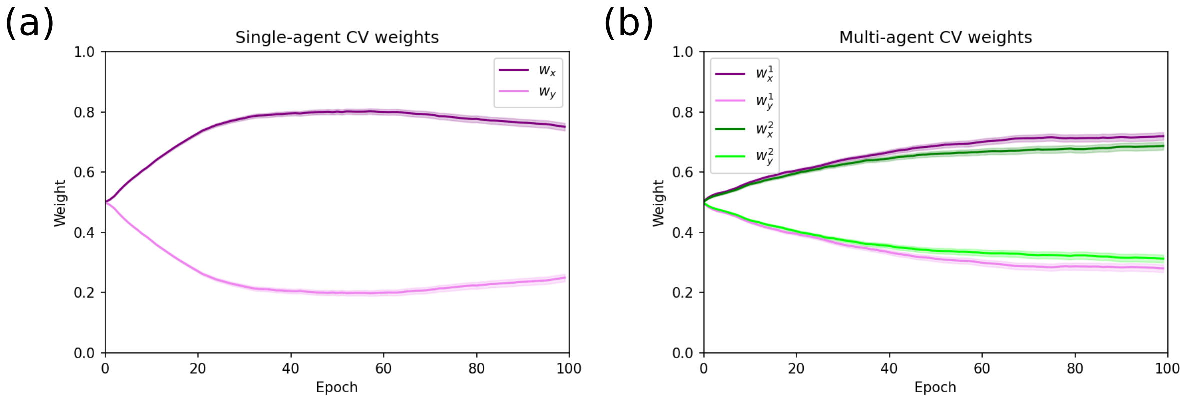


Figure S1: Collective variable weights for the symmetric cross potential trials employing (a) SA REAP and (b) MA REAP. $w_\theta^a$ represents the weight that agent $a$ assigns to variable $\theta$. The curves show means ± 95% confidence intervals over all replicates. Since the agents are initially placed at the extremes of the horizontal arms of the cross, we observe that $w_x > w_y$ (for both agents). This trend continues for the entire run due to the fact that, even after discovering the vertical arms, exploration along the $x$-axis continues being rewarding.

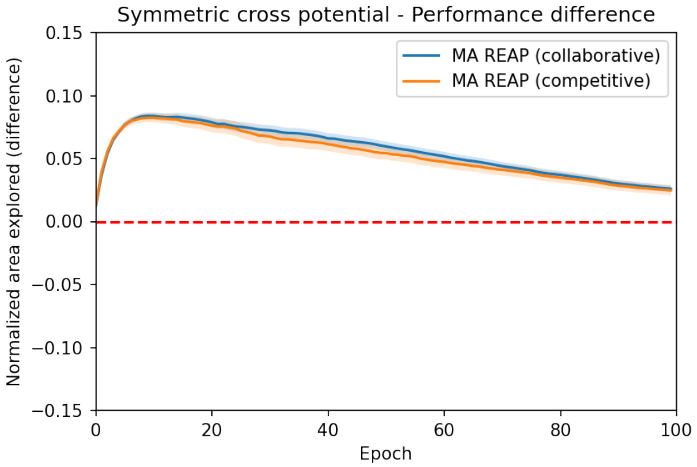


Figure S2: Comparison between MA REAP utilizing collaborative and competitive rewards vs. Least Counts adaptive sampling. Curves show means ±95% confidence intervals. No significant difference is observed between the two MA REAP runs employing different reward combination regimes.

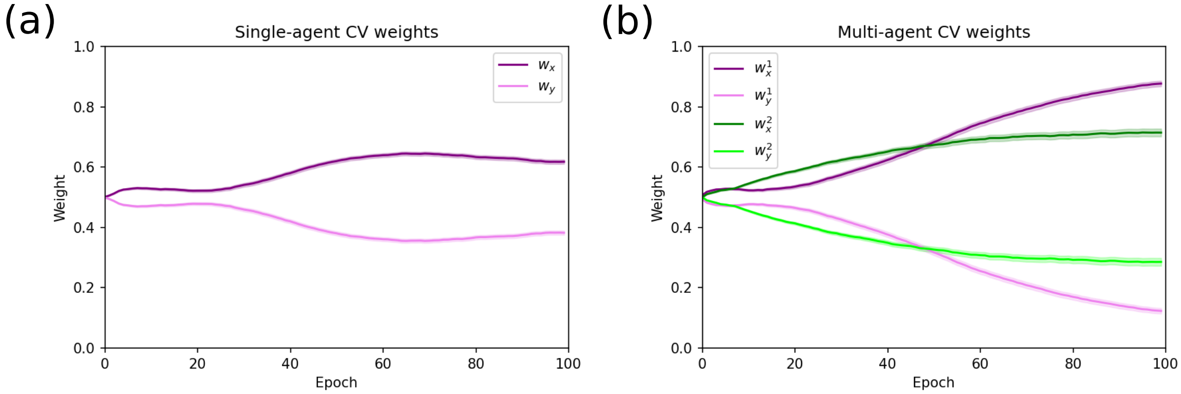


Figure S3: Collective variable weights for the asymmetric cross potential trials employing (a) SA REAP and (b) MA REAP. $w_\theta^a$ represents the weight that agent $a$ assigns to variable $\theta$. The curves show means $\pm$ 95% confidence intervals over all replicates. Since the agents are initially placed at the extremes of the horizontal arms of the cross, we observe that $w_x > w_y$ (for both agents). This trend continues for the entire run due to the fact that, even after discovering the vertical arms, exploration along the $x$-axis continues being rewarding. We observe that $w_x^1 \approx 1$ because agent 1 is constrained to the horizontal arm due to the deep minimum it is initially placed on.

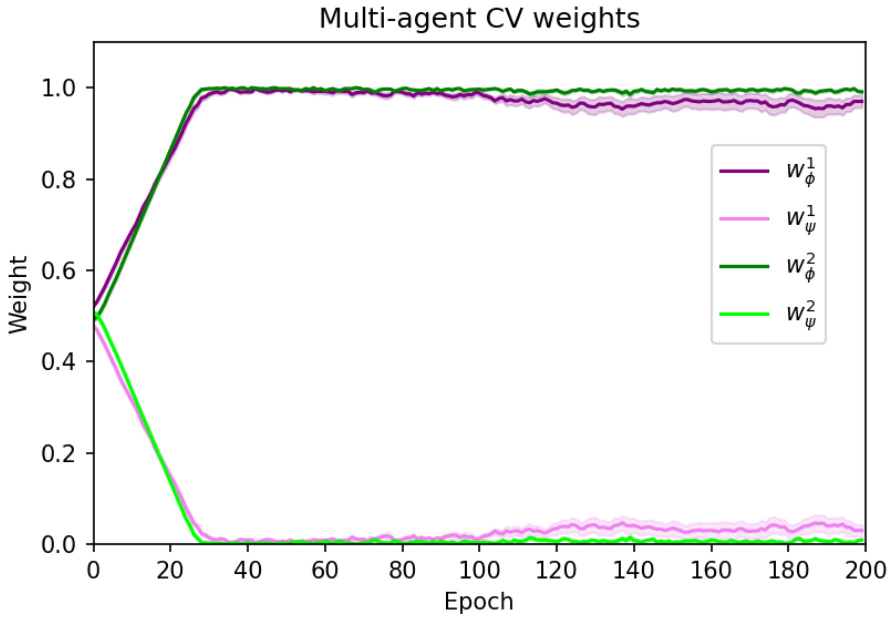


Figure S4: Collective variable weights for the alanine dipeptide trials. $w_\theta^a$ represents the weight that agent $a$ assigns to variable $\theta$. $\phi$ and $\psi$ are the dihedral angles as traditionally defined. The curves show means $\pm$ 95% confidence intervals over all replicates. We observe that $w_\phi \gg w_\psi$ for all agents because the $\psi$ dihedral can be readily explored by both agents and the discovery of the states with $\phi > 0$ results in high rewards.

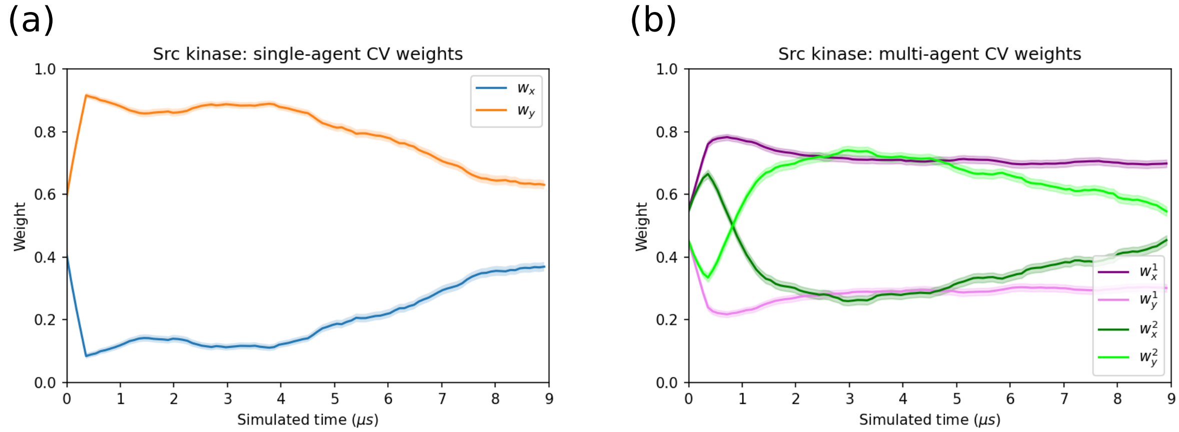


Figure S5: Collective variable weights for the Src kinase trials employing (a) SA REAP and (b) MA REAP. $w_\theta^a$ represents the weight that agent $a$ assigns to variable $\theta$. $x$ represents the RMSD of the A-loop and $y$ is the distance between K295 and E310. The curves show means $\pm$ 95% confidence intervals over all replicates. In (b), agents 1 and 2 start at the inactive and active states, respectively. We observe $w_x^1 > w_y^1$ because it is more rewarding for agent 1 to explore along the RMSD CV (see Figure 4a in Main Text). On the other hand, agent 2 learns after 1 $\mu$s that sampling along the distance variable is more favorable. This delay in reaching $w_y^2 > w_x^2$ is caused by the discovery of states in the region where $y \approx 0.4$ nm and $x \in [0.7, 1]$ nm, which misleads the agent by suggesting that exploration along $x$ is more rewarding than along $y$ in that region of the landscape.

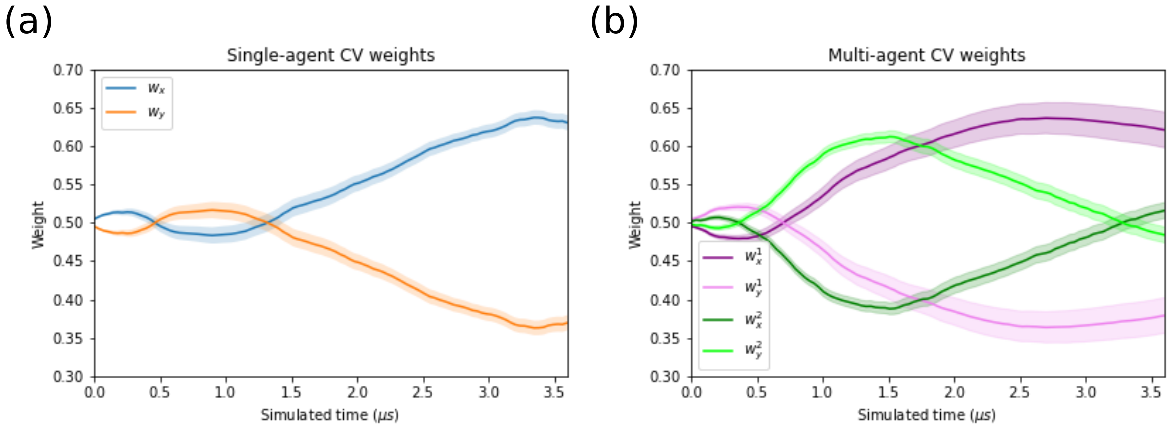


Figure S6: Collective variable weights for the OsSWEET2b trials employing (a) SA REAP and (b) MA REAP. $w_\theta^a$ represents the weight that agent $a$ assigns to variable $\theta$. $x$ represents the periplasmic distance (distance between D190 and R70) and $y$ is the cytoplasmic distance distance between F165 and F43). The curves show means ± 95% confidence intervals over all replicates. In (b), agents 1 and 2 start at the inward facing and outward facing states, respectively.

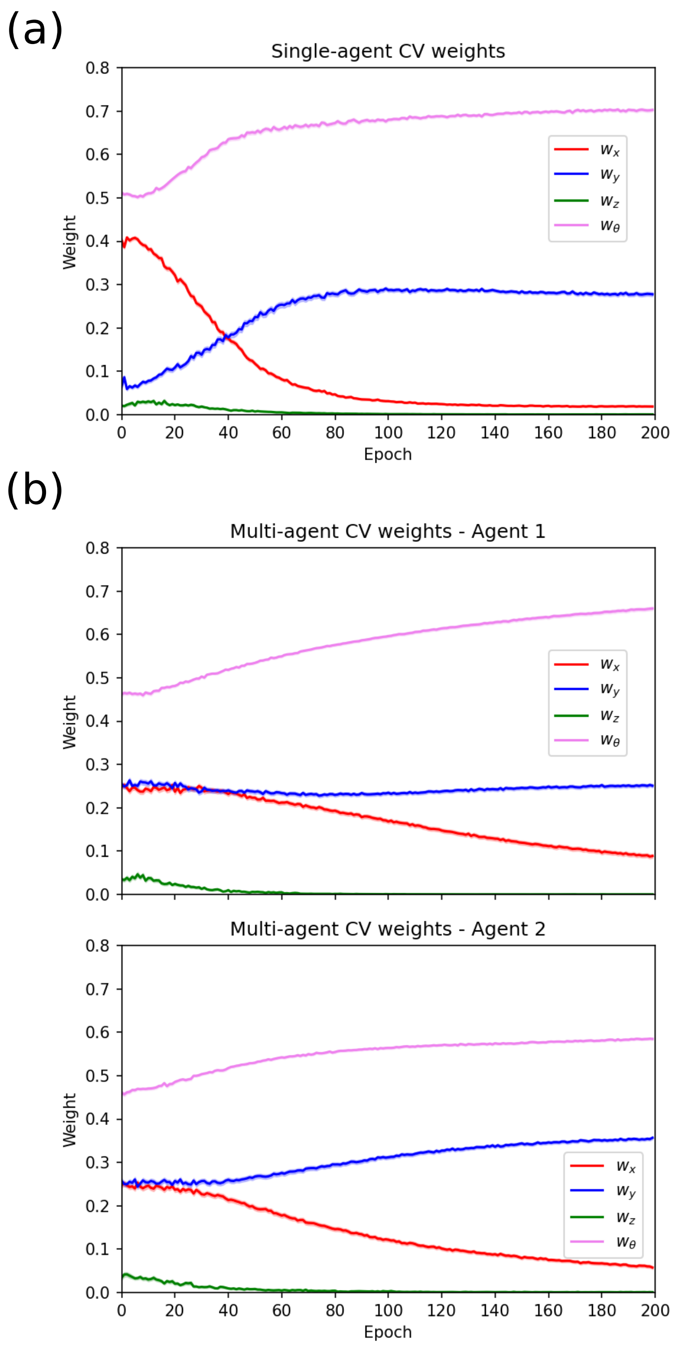


Figure S7: CV weights derived using the original (a) and multi-agent (b) implementations of TSLC. The curves show means ± 95% confidence intervals (imperceptible) over all replicates. In all cases, the angle CV is assigned the highest weight because the gradient of this variable possesses the largest projection on the local tangent space across clusters. A large $w_\theta$ promotes the exploration of the toroidal potential.

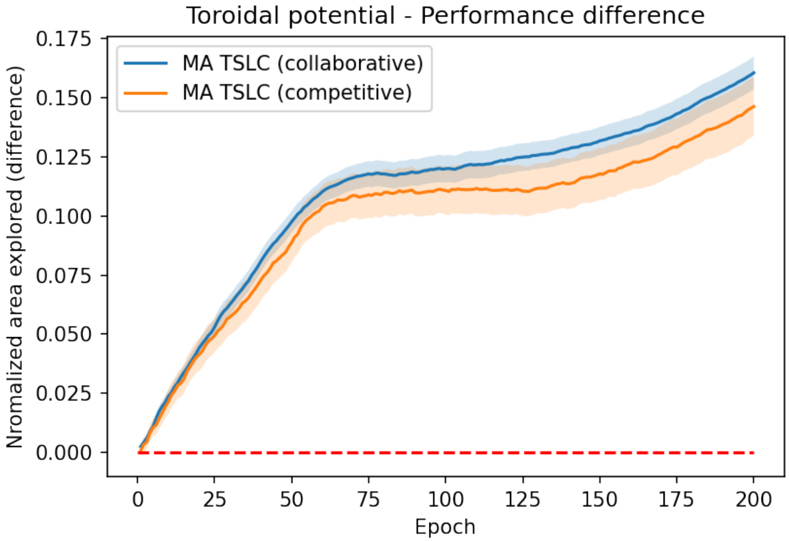


Figure S8: Comparison between MA TSLC utilizing collaborative and competitive rewards vs. SA TSLC. Curves show means ±95% confidence intervals. No significant difference is observed between the two MA TSLC runs employing different reward combination regimes.

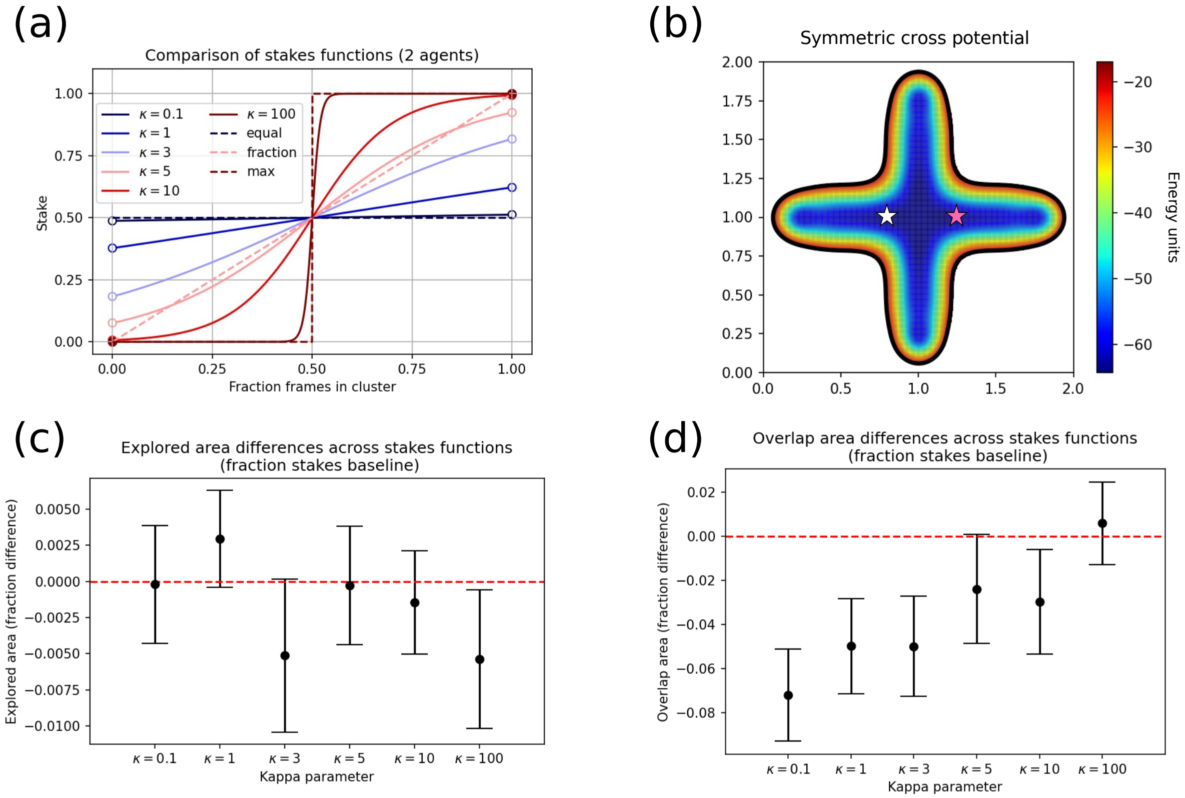


Figure S9: Stakes functions comparison on the symmetric cross potential. (a) Stakes functions for the two-agent case. Solid lines correspond to logistic stakes with $\kappa$ values indicated in the legend. These functions present discontinuities at the ends because they must evaluate to 0 or 1 at the extrema. Dashed lines correspond to equal, fraction, and max functions. (b) Symmetric cross potential used in the comparison. Starting points (marked by stars) are identical to those used in the other comparisons. (c) Difference in explored area. The error bars represent 95% confidence intervals. Result is statistically significant if the confidence interval does not cross zero. (d) Difference in area overlap. The error bars represent 95% confidence intervals. Result is statistically significant if the confidence interval does not cross zero.

(S2) Buenfil, J.; Koelle, S. J.; Meila, M. Tangent Space Least Adaptive Clustering. ICML 2021 Workshop on Unsupervised Reinforcement Learning. 2021.

(S3) Harvey, M. J.; Giupponi, G.; Fabritiis, G. D. *J. Chem. Theory Comput.* **2009**, *5*, 1632–1639.

(S4) Harrigan, M. P.; Sultan, M. M.; Hernández, C. X.; Husic, B. E.; Eastman, P.; Schwantes, C. R.; Beauchamp, K. A.; McGibbon, R. T.; Pande, V. S. *Biophys. J.* **2017**, *112*, 10–15.

(S5) Shamsi, Z.; Cheng, K. J.; Shukla, D. *J. Phys. Chem. B* **2018**, *122*, 8386–8395.

(S6) Pérez, A.; Herrera-Nieto, P.; Doerr, S.; Fabritiis, G. D. *J. Chem. Theory Comput.* **2020**, *16*, 4685–4693.

(S7) Shukla, D.; Meng, Y.; Roux, B.; Pande, V. S. *Nat. Commun.* **2014**, *5*, 3397.

(S8) Selvam, B.; Yu, Y.-C.; Chen, L.-Q.; Shukla, D. *ACS Cent. Sci.* **2019**, *5*, 1085–1096.

(S9) Scherer, M. K.; Trendelkamp-Schroer, B.; Paul, F.; Pérez-Hernández, G.; Hoffmann, M.; Plattner, N.; Wehmeyer, C.; Prinz, J.-H.; Noé, F. *J. Chem. Theory Comput.* **2015**, *11*, 5525–5542.
